# Speciation in sympatry with ongoing secondary gene flow and an olfactory trigger in a radiation of Cameroon cichlids

**DOI:** 10.1101/229864

**Authors:** Jelmer W. Poelstra, Emilie J. Richards, Christopher H. Martin

## Abstract

Whether speciation can happen in the absence of geographical barriers and if so, under which conditions, is a fundamental question in our understanding of the evolution of new species. Among candidates for sympatric speciation, Cameroon crater lake cichlid radiations have been considered the most compelling. However, it was recently shown that a more complex scenario than a single colonization followed by isolation underlies these radiations. Here, we perform a detailed investigation of the speciation history of a radiation of *Coptodon* cichlids from Lake Ejagham using whole-genome sequencing data. The existence of the Lake Ejagham *Coptodon* radiation is remarkable since this 0.5 km^2^ lake offers limited scope for divergence across a shallow depth gradient, disruptive selection is weak, the species are sexually monochromatic, yet assortative mating is strong. We infer that Lake Ejagham was colonized by *Coptodon* cichlids almost as soon as it came into existence 9,000 years ago, yet speciation events occurred only in the last 1,000-2,000 years. We show that secondary gene flow from a nearby riverine species has been ongoing, into ancestral as well as extant Lake Ejagham lineages, and we identify and date river-to-lake admixture blocks. One of these contains a cluster of olfactory receptor genes that introgressed close to the time of the first speciation event and coincides with a higher overall rate of admixture into the recipient lineages. Olfactory signaling is a key component of mate choice and species recognition in cichlids. A functional role for this introgression event is consistent with previous findings that assortative mating appears much stronger than ecological divergence in Ejagham *Coptodon.* We conclude that speciation in this radiation took place in sympatry, yet may have benefited from ongoing riverine gene flow.

**Author Summary:** Despite an active search for empirical examples and much theoretical work, sympatric speciation remains one of the most controversial ideas in evolutionary biology. While a host of examples have been described in the last few decades, more recent results have shown that several of the most convincing systems have not evolved in complete isolation from allopatric populations after all. By itself, documenting the occurrence of secondary gene flow is not sufficient to reject the hypothesis of sympatric speciation, since speciation can still be considered sympatric if gene flow did not contribute significantly to the build-up of reproductive isolation. One way forward is to use genomic data to infer where, when and into which lineages gene flow occurred, and identify the regions of the genome that experienced admixture. In this study, we use whole-genome sequencing to examine one of the cichlid radiations from a small isolated Cameroon lake, which have long been the flagship example of sympatric speciation. We show that gene flow from a riverine species into the lake has been ongoing during the history of the radiation. In line with this, we infer that the lake was colonized very soon after it was formed, and argue that Lake Ejagham is not as isolated as previously assumed. The magnitude of secondary gene flow was relatively even across Lake Ejagham lineages, yet with some evidence for differential admixture, most notably before the first speciation event into the *C. deckerti* and *C. ejagham* lineage. Among the sequences that were introgressed into this lineage is a cluster of olfactory receptor genes, which may have facilitated speciation by promoting sexual isolation between incipient species, consistent with previous findings that sexual isolation appears to be stronger than ecological isolation in Ejagham *Coptodon*. We conclude that speciation in this radiation took place in sympatry, yet may have benefited from ongoing riverine gene flow.

## Introduction

Speciation in the absence of geographic barriers is a powerful demonstration that divergent selection can overcome the homogenizing effects of gene flow and recombination (Arnegard and Kondrashov, 2004, 2004; Coyne and Orr, 2004; Turelli et al., 2001). While it was long thought that sympatric speciation was very unlikely to take place in nature, the last 25 years have seen a proliferation of empirical examples as well as theoretical models that support its plausibility (Barluenga et al., 2006; Berlocher and Feder, 2002; Bolnick and Fitzpatrick, 2007; Hadid et al., 2013, 2014, Kautt et al., 2016a, 2016b; Malinsky et al., 2015; Savolainen et al., 2006; Sorenson et al., 2003).

However, it is exceptionally hard to demonstrate that speciation has been sympatric in any given empirical case. One of the most challenging criteria is that there has been no historical phase of geographic isolation (Coyne and Orr, 2004). This can be ruled out most compellingly in cases where multiple endemic species are found in environments that are (i) small and homogeneous, such that geographic isolation within the environment is unlikely, and are (ii) severely isolated, such that a single colonization likely produced the lineage that eventually diversified. Early molecular studies used a single locus or limited genomic data to establish monophyly of sympatric species in isolated environments such as crater lakes (Barluenga et al., 2006; Schliewen et al., 1994) and oceanic islands (Savolainen et al., 2006). Genome-wide sequencing data can now be used to rigorously test whether or not extant species contain ancestry from secondary gene flow into the environment. Strikingly, evidence for such ancestry has recently been found in all seven crater lake cichlid radiations examined so far (Kautt et al., 2016b; Malinsky et al., 2015; Martin et al., 2015a). However, whereas a complete lack of secondary gene flow would rule out a role for geographic isolation outside of the focal environment, the presence of secondary gene flow does not exclude the possibility of sympatric speciation.

If secondary gene flow into a pair or radiation of sympatric species has taken place, a key question is whether secondary gene flow played a role in the speciation process. Speciation would still be functionally sympatric if genetic variation introduced by secondary gene flow did not contribute to speciation (Martin et al., 2015a). Secondary gene flow could even counteract speciation in the focal environment, for instance via hybridization with both incipient species during a speciation event. On the other hand, there are several ways in which secondary gene flow may be a key part of speciation (Kautt et al., 2016b; Martin et al., 2015a). For instance, secondary colonization may involve a partially reproductively isolated population, in which case any resulting speciation event would have a crucial allopatric phase. Second, the introduction of novel genetic variation and novel allelic combinations may promote speciation more generally; for example, through the formation of a hybrid swarm (Seehausen, 2004).

Establishing or rejecting a causal role of secondary gene flow in speciation can be very difficult in any particular case, yet a first step is to examine the timing, extent, and identity of the donor and recipient populations. A role for secondary gene flow would be supported if divergence rapidly followed a discrete admixture event (Kautt et al., 2016b); whereas, if gene flow took place only after the onset of divergence, such a role would seem unlikely. Genomic data can also be used to identify segments of the genome that have experienced admixture and to examine whether these contain genes that may have been important in speciation (Lamichhaney et al., 2015; Meier et al., 2017; Richards and Martin, 2017).

Four radiations of cichlids in three isolated lakes in Cameroon (Schliewen and Klee, 2004; Schliewen et al., 2001, 1994) are one of the most widely accepted examples of sympatric speciation. Two of the lakes are crater lakes, while the third, Lake Ejagham, is now suspected to be the result of a meteor impact (Stager et al., 2017). Given their small size and uniform topology, geographic isolation within these lakes is unlikely (Schliewen et al., 1994). Moreover, species within the radiations were shown to be monophyletic relative to riverine outgroups based on mtDNA (for all four of the radiations, Schliewen et al., 1994) and AFLPs (for one radiation, Schliewen and Klee, 2004), which was interpreted to mean that each radiation is derived from a single colonization. However, using RAD-seq data, Martin et al. (2015a) recently found evidence for secondary admixture with nearby riverine populations in all four radiations.

Despite being the second smallest (0.49 km^2^) and one of the youngest lakes (ca. 9,000 ka, Stager et al., 2017) containing endemic cichlids, Lake Ejagham contains two independent endemic cichlid radiations, a two-species radiation of *Sarotherodon* cichlids (Neumann, 2011) and a four-species radiation of *Coptodon* cichlids (Dunz and Schliewen, 2010). The existence of these radiations is all the more remarkable given that they are an exception to the two best predictors of endemic radiation in African cichlids: lake depth and sexual dichromatism (Wagner et al., 2012). Sympatric cichlid species pairs are commonly distributed over a large depth gradient (Kautt et al., 2016a, 2016b; Malinsky et al., 2015), yet Lake Ejagham is shallow (maximum depth of 18 m, Schliewen et al., 2001), and at least three species are completely interspersed in the same depth range (Martin, 2013). While no sexual dichromatism occurs among Ejagham *Coptodon*, all species differ most strongly in sexual rather than ecological characters (Martin, 2013), and strong assortative mating appears to be a more significant force than weak disruptive selection (Martin, 2012), which is noteworthy since speciation in Cameroon lakes is generally considered to be ecologically driven (Coyne and Orr, 2004).

Some of the clearest evidence for admixture in (Martin et al., 2015a) came from the three species of Ejagham *Coptodon* that were examined. The occurrence of secondary gene flow from riverine populations could be a key piece in the puzzling occurrence of the Lake Ejagham radiations, and may have initiated speciation despite limited disruptive ecological selection. Here, we use whole-genome sequencing of three species of Ejagham *Coptodon* and two riverine outgroups to provide a comprehensive picture of the history of secondary gene flow and its riverine sources, and identify admixed portions of the genome.

## Results

### Phylogeny of the Lake Ejagham Coptodon radiation

As a first step in revealing the speciation history of the Lake Ejagham *Coptodon* radiation (hereafter: Ejagham radiation), we took several approaches to infer the phylogenetic relationships among the three Lake Ejagham species *C. deckerti, C. ejagham* and *C. fusiforme*, as well as two closely related riverine species from the neighboring Cross River drainage, *C. guineensis* and *C. sp. Mamfé* (Fig 1), *C. kottae*, a Cameroon crater lake endemic that did not diversify in situ, and the much more distantly related *Sarotherodon galilaeus.*

**Fig 1.**
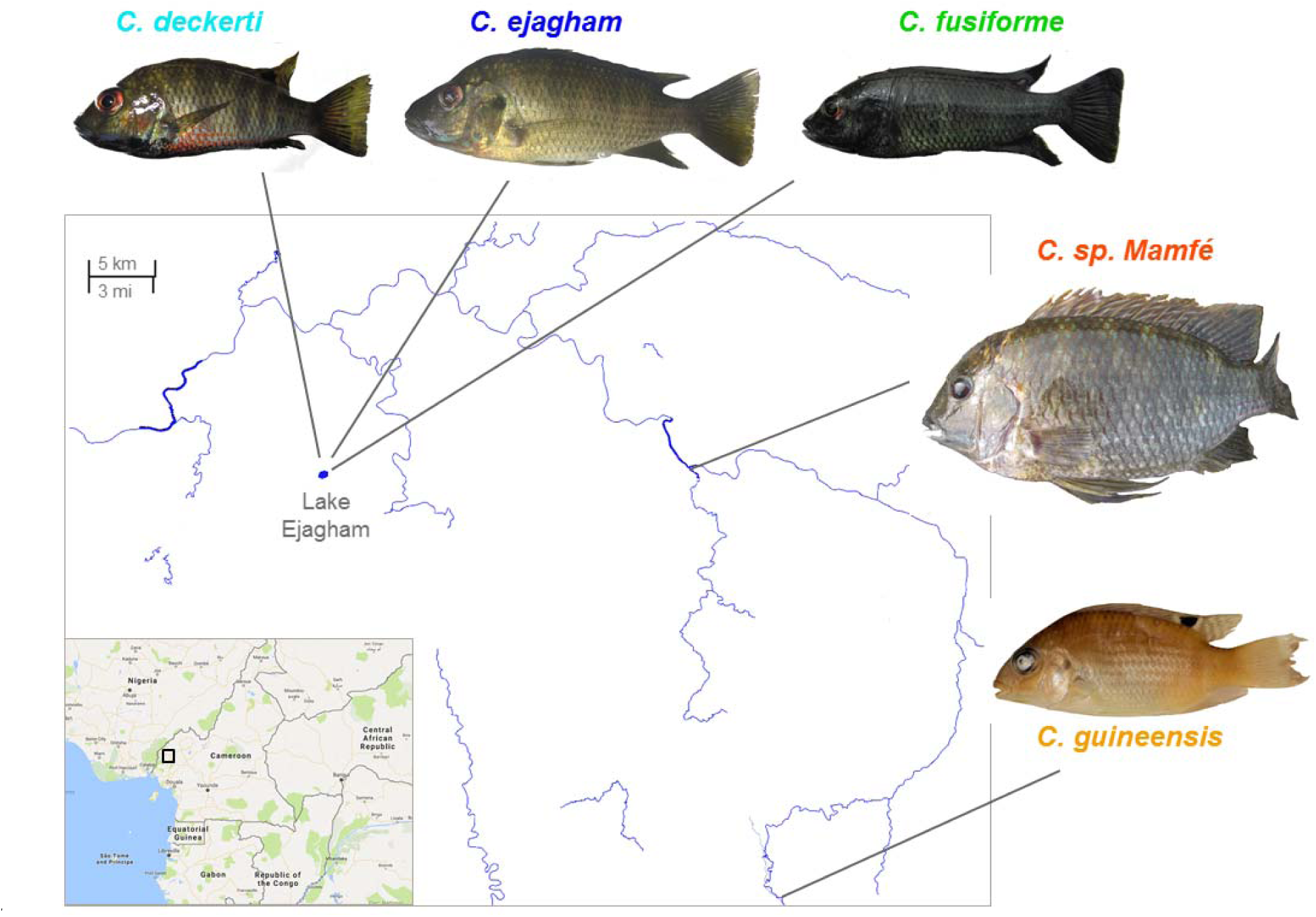
Lake Ejagham and its surrounding rivers in western Cameroon. The focal species in this study are shown: three species of Lake Ejagham *Coptodon* and two closely related riverine species. As outgroups, we used *C. kottae*, a Cameroon crater lake endemic species that did not diversify, and *Sarotherodon galilaeus.*

Maximum likelihood (ML) trees based on concatenated genome-wide SNPs using RaxML with any of three outgroup configurations (only *C. kottae* / only *S. galilaeus* / both species) resulted in monophyly of Lake Ejagham species and a sister relationship between *C. deckerti* and *C. ejagham* with 100% bootstrap support (Fig 2A). However, inferences on whether one of the two riverine species is more closely related to Ejagham *Coptodon*, or the two are sister species, differed among outgroup configurations (Fig S1). To further investigate the relationships among the two riverine species relative to Ejagham *Coptodon*, we constructed species trees based on 100 kb gene trees. Species trees based on rooted gene trees using ML and the Minimize Deep Coalescence (MDC) criterion in Phylonet, as well as a species tree based on unrooted gene trees using ASTRAL, all indicated monophyly of the Ejagham radiation, and a sister relationship between *C. deckerti* and *C. ejagham* (Fig S2).

**Fig 2.**
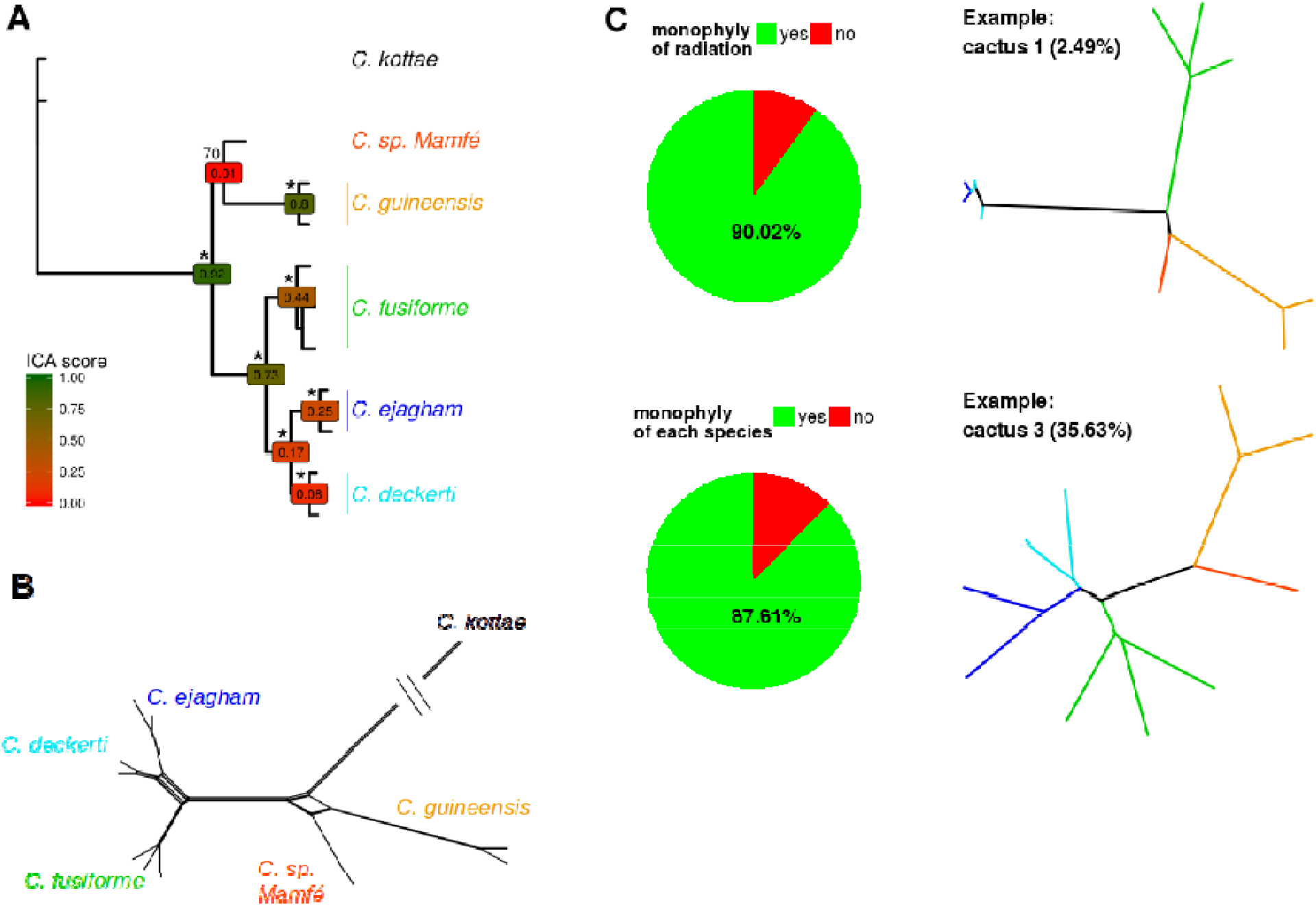
Support for monophyly of the Lake Ejagham *Coptodon* radiation across the genome. **(A)** Maximum likelihood tree based on concatenated SNPs across the genome, with bootstrap support (* = 100% support), and ICA (Internode Confidence All) values based on ML gene trees for 100kb windows. Support for the sister relationship between the riverine species *C. sp. Mamfé* and *C. guineensis* is much lower than that for the monophyly of the three lake Ejagham species, *C. fusiforme, C. ejagham*, and *C. deckerti*. **(B)** A phylogenetic network shows limited conflict along the branch leading to lake Ejagham species and a rather clearly resolved topology within the radiation. In line with results from panel A, more conflict is observed around the divergence of *C. sp. Mamfé* and *C. guineensis*. **(C)** Local phylogenies (Saguaro “cacti”) indicate that along most of the genome, the Ejagham *Coptodon* clade (top) is monophyletic and that individuals within the clade cluster by species (bottom).

We used two methods to more explicitly examine the prevalence of discordant phylogenetic patterns. In keeping with the results from phylogenetic trees, a phylogenetic network based on genome-wide SNPs produced by Splitstree showed limited discordance along the branch to the Ejagham *Coptodon* ancestor, with higher levels of discordance along the branch to the *C. deckerti - C. ejagham* ancestor and especially near the divergence of the riverine species (Fig 2B). Second, phylogenetic relationships along local segments of the genome grouped by the machine-learning approach *Saguaro* into 30 unrooted trees (“cacti”) indicate that in 90.02% of the genome, Ejagham *Coptodon* and the two riverine species each form exclusive clades (Fig 2C, S3 Fig, S7 Table). Similarly, in 87.61% of the genome, individuals in each of the three Ejagham species grouped monophyletically (Fig 2C, S7 Table).

### Genome-wide tests of admixture suggest ongoing gene flow from C. sp. Mamfé

To further investigate admixture between riverine and Lake Ejagham taxa, we first used genome-wide formal tests of admixture. Genome-wide D-statistics in configurations that test for admixture between one of the two riverine species and an Ejagham *Coptodon* species, repeated for each Ejagham species, all indicate admixture between *C. sp. Mamfé* and Ejagham *Coptodon* (Fig 3A, top three bars). Values of *D* were very similar (0.1578 - 0.1594) across the three Ejagham species, indicating similar levels of admixture from *C. sp. Mamfé*. This suggests that admixture may have predominantly taken place prior to diversification within Lake Ejagham.

**Fig 3.**
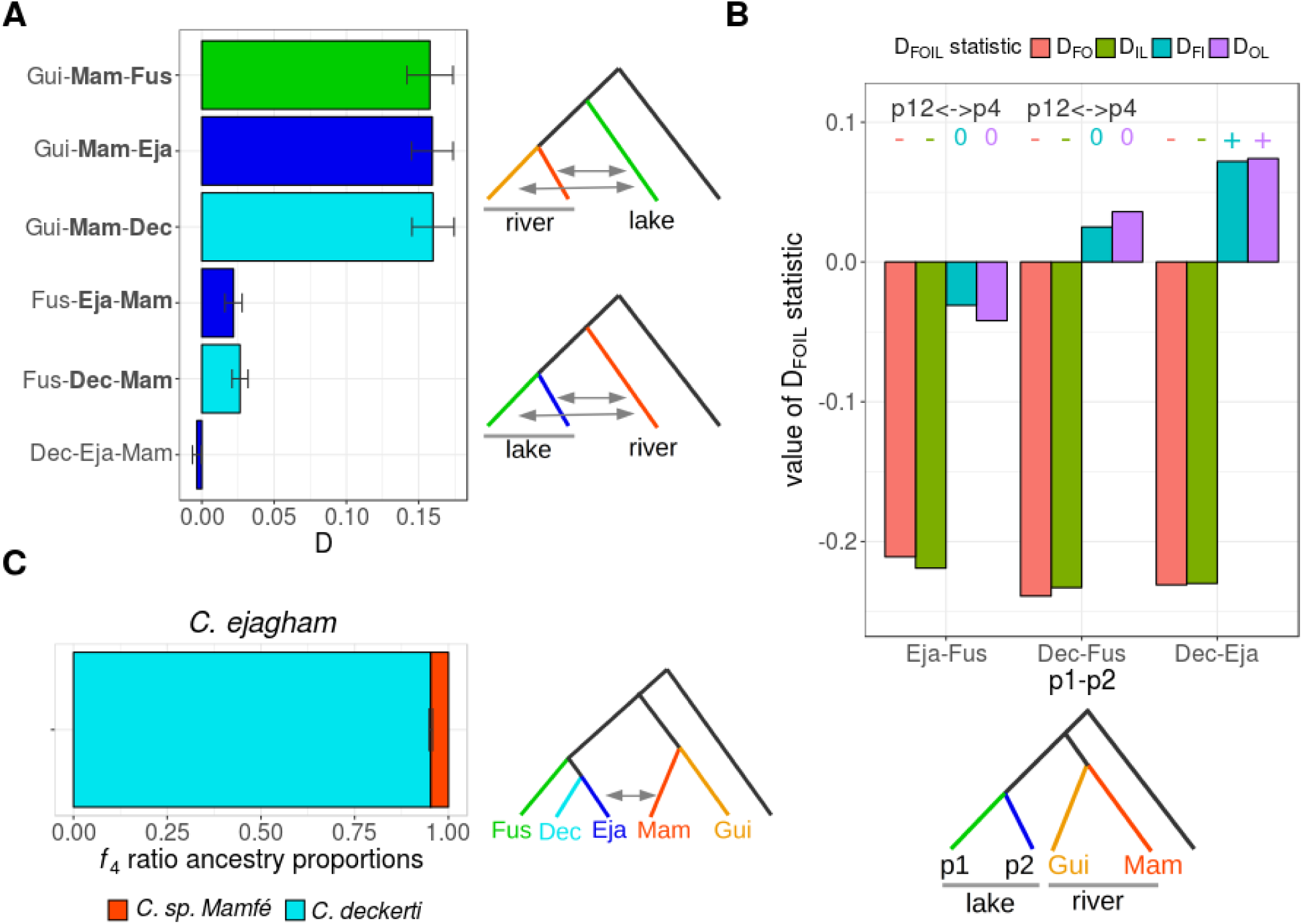
Genome-wide admixture statistics suggest secondary riverine gene flow from *C. sp. Mamfé*. (**A**) D-statistics for several ingroup triplets indicate that all three Ejagham *Coptodon* species (”Fus”: *C. fusiforme*, “Eja”: *C. ejagham*, “Dec”: *C. deckerti*) experienced admixture with *C. sp. Mamfé* (“Mam”), at similar levels relative to *C. guineensis* (“Gui”), as shown by the top three bars. The lower three bars show the much weaker evidence for differential *C. sp. Mamfé* admixture among Ejagham *Coptodon* species. Species between which admixture is inferred (significant D-statistics) are denoted in bold. (**B**) D_FOIL_ statistics for the three combinations of two Ejagham *Coptodon* species show a preponderance of ancestral gene flow with *C. sp. Mamfé*. Negative D_FO_ and D_IL_ in combination with non-significant D_FI_ and D_OL_ statistics, as for the first two comparisons, indicate ancestral gene flow, while the pattern for the third combination does not have a straightforward interpretation, although it is qualitatively similar to the first two comparisons. (**C**) An *f_4_*-ratio test for differential *C. sp. Mamfé* admixture between *C. ejagham* and *C. deckerti* indicates that *C. ejagham* has experienced 4.7% additional admixture from *C. sp. Mamfé*.

We tested this interpretation using five-taxon D_FOIL_ statistics (Fig 3B). D_FOIL_ statistics take advantage of derived allele frequency patterns in a symmetric phylogeny across two pairs of populations with dissimilar coalescence times. The combination of signs (positive, negative, or zero) across four D_FOIL_ statistics can distinguish between admixture along terminal branches and admixture with the ancestral population of the most recently diverged population pair. Here, we repeated the test with each of three possible pairs of Lake Ejagham species as P1 and P2, and as P3 and P4 the pair of riverine species, which diverged prior to the Ejagham species (see next section). D_FOIL_ statistics using both pairs of Lake Ejagham taxa that involve *C. fusiforme* indicated a pattern of admixture between *C. sp. Mamfé* and the Lake Ejagham ancestor (Fig 3B, left). D_FOIL_ statistics are designed to uncover a single admixture pattern, such that multiple instances of gene flow may lead to a combination of signs across D_FOIL_ statistics without a straightforward interpretation, which may explain the pattern observed for the comparison with *C. deckerti* and *C. ejagham* as P1 and P2 (Fig 3B, right).

Consistent with more complex patterns of admixture, D-statistics for comparisons that explicitly test for differential admixture between Ejagham species with *C. sp. Mamfé* indicate that *C. ejagham* and *C. deckerti* experienced slightly higher levels of admixture than *C. fusiforme* after their divergence (Fig 3A, bottom bars). Furthermore, an *f_4_*-ratio test suggests that 4.7% of *C. ejagham* ancestry derives from admixture with *C. sp. Mamfé* during or after its divergence from *C. deckerti* (Fig 3C), but it should be noted that D-statistics did not indicate differential admixture for this comparison (Fig 3A, bottom bar). Overall, we infer that differential gene flow from *C. sp. Mamfé* into the three Ejagham species has been relatively minor in comparison to gene flow shared among the species. This difference in magnitude can be clearly seen in Fig 3A, where the upper three bars represent shared gene flow and the lower three bars differential gene flow to Ejagham species.

### A detailed reconstruction of the demographic speciation history of the Ejagham radiation

To infer post-divergence rates of gene flow, divergence times, and population sizes among the extant and ancestral Lake Ejagham lineages and the two riverine species, we used the Generalized Phylogenetic Coalescent Sampler (G-PhoCS), providing the species tree topology inferred above. Gene flow rates in G-PhoCS can be estimated using specific “migration bands” between any two lineages that overlap in time. We focused on migration bands that had a riverine lineage as the source population and an extant or ancestral Lake Ejagham lineage as the target population. We first inferred rates in models with single migration bands, and next combined significant migration bands in models with multiple migration bands. While models with all migration bands performed more poorly due to the high number of parameters (see Methods), models with single migration bands may be prone to overestimation of that specific migration rate. We therefore also ran models with an intermediate number of migration bands (either to all three extant Ejagham species or to both ancestral lineages), and present results for all of these different models in Fig 4, and Table 1. Divergence times and population sizes mentioned below represent only those from models with all significant migration bands.

**Fig 4.**
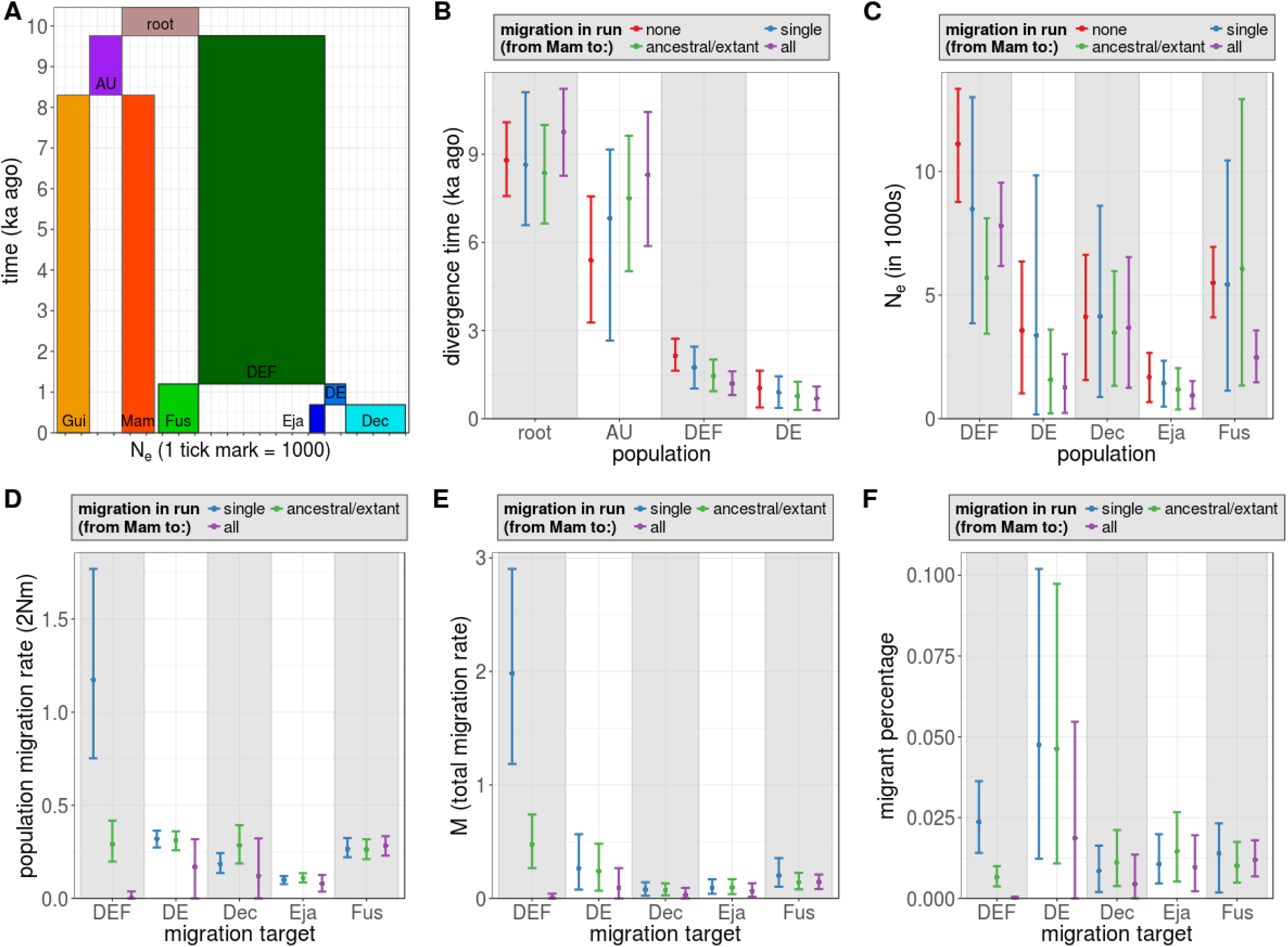
A comprehensive picture of the demographic speciation history of Coptodon *Ejagham*. (**A**) Overview of the divergence times and population sizes inferred by G-PhoCS. Box widths (x-axis) correspond to population sizes only for Lake Ejagham lineages: *C. deckerti*(“Dec”), C. ejagham (“Eja”), C. fusiforme (“Fus”), the ancestor of Dec and Eja (“DE”), and the ancestral Ejagham lineage (“DEF”). (**B-F**) Estimates of divergence times (B), population sizes (C), and migration rates (D-F) across runs with varying migration bands from *C. sp. Mamfé* to lake lineages: “none”, “single”, “ancestral/current”, and “all” indicate that individual runs estimated zero, one, several (either to the two ancestral lineages, DE and DEF, or to the three extant species), or all possible migration bands, respectively.

**Table 1.** **Summary of G-PhoCS parameter estimates.** Divergence time τ represents the estimated time that the named lineage split into its daughter lineage (see Fig, 4A). All migration rates are from migration from *C. sp. Mamfé* to Lake Ejagham lineages. Parameter estimates are given separately for runs with no migration (“none”), with a single migration band (“single”), with migration bands to either both ancestral or all three extant lineages (“anc/ext”), or to all Lake Ejagham lineages. “τ” – divergence time; “2Nm” – population migration rate; “M (total)” – total migration rate; “% migrants” – percentage of migrants received in each generation; “AU” – ancestor of *C. sp. Mamfé* and *C. guineensis;* “DEF” – ancestor of all three lake Ejagham species; “DE” – ancestor of *C. deckerti*and *C. ejagham; Dec* – “C. deckerti”; “Eja” – *C. Ejagham;* “Fus” – *C. fusiforme*.

| parameter | lineage | mean<br>single | mean<br>anc/ext | mean<br>all | mean<br>none | 95% HPD<br>single | 95% HPD<br>anc/ext | 95% HPD<br>all | 95% HPD<br>none |
| --- | --- | --- | --- | --- | --- | --- | --- | --- | --- |
| $\tau$ | root | 8,649 | 8,369 | 9,760 | 8,803 | 6,587-11,112 | 6,647-10,001 | 8,267-11,229 | 7,579-10,090 |
| $\tau$ | AU | 6,823 | 7,498 | 8,298 | 5,393 | 2,658-9,163 | 5,024-9,631 | 5,880-10,438 | 3,275-7,571 |
| $\tau$ | DEF | 1,740 | 1,454 | 1,205 | 2,150 | 1,027-2,451 | 936-2,015 | 806-1,616 | 1,633-2,721 |
| $\tau$ | DE | 892 | 778 | 689 | 1,049 | 365-1,443 | 300-1,259 | 291-1,096 | 3,83-1,635 |
| $N_e$ | DEF | 8,482 | 5,714 | 7,794 | 11,121 | 3,857-13,001 | 3,435-8,109 | 6,175-9,545 | 8,768-13,343 |
| $N_e$ | DE | 3,373 | 1,589 | 1,288 | 3,574 | 171-9,846 | 216-3,608 | 235-2,613 | 1,025-6,358 |
| $N_e$ | Dec | 4,133 | 3,500 | 3,681 | 4,128 | 874-8,615 | 1,328-5,967 | 1,250-6,539 | 1,566-6,625 |
| $N_e$ | Eja | 1,425 | 1,180 | 933 | 1,684 | 489-2,343 | 371-2,044 | 406-1,525 | 670-2,662 |
| $N_e$ | Fus | 5,432 | 6,069 | 2,474 | 5,488 | 1,131-10,444 | 1,342-12,925 | 1,469-3,572 | 4,100-6,946 |
| 2Nm | DEF | 1.18 | 0.29 | 0.01 | NA | 0.75-1.77 | 0.2-0.42 | 0-0.04 | NA |
| 2Nm | DE | 0.32 | 0.31 | 0.17 | NA | 0.27-0.36 | 0.26-0.36 | 0-0.32 | NA |
| 2Nm | Dec | 0.19 | 0.28 | 0.12 | NA | 0.14-0.24 | 0.19-0.39 | 0-0.32 | NA |
| 2Nm | Eja | 0.10 | 0.11 | 0.08 | NA | 0.08-0.12 | 0.09-0.14 | 0.04-0.13 | NA |
| 2Nm | Fus | 0.27 | 0.26 | 0.28 | NA | 0.22-0.32 | 0.21-0.32 | 0.23-0.33 | NA |
| M (total) | DEF | 1.98 | 0.48 | 0.01 | NA | 1.18-2.9 | 0.27-0.74 | 0-0.04 | NA |
| M (total) | DE | 0.27 | 0.24 | 0.09 | NA | 0.08-0.57 | 0.07-0.48 | 0-0.27 | NA |
| M (total) | Dec | 0.08 | 0.07 | 0.03 | NA | 0.02-0.14 | 0.03-0.13 | 0-0.09 | NA |
| M (total) | Eja | 0.09 | 0.09 | 0.07 | NA | 0.04-0.17 | 0.04-0.17 | 0.01-0.13 | NA |
| M (total) | Fus | 0.20 | 0.14 | 0.14 | NA | 0.1-0.35 | 0.08-0.23 | 0.08-0.21 | NA |
| % migrants | DEF | 1.39e-4 | 5.07e-5 | 7.03e-7 | NA | 8.9e-5-2.1e-4 | 3.5e-5-7.3e-5 | 0-4.8e-6 | NA |
| % migrants | DE | 9.47e-5 | 1.96e-4 | 1.33e-4 | NA | 8.1e-5-1.1e-4 | 1.6e-4-2.3e-4 | 0-2.5e-4 | NA |
| % migrants | Dec | 4.52e-5 | 8.11e-5 | 3.33e-5 | NA | 3.3e-5-5.9e-5 | 5.4e-5-1.1e-4 | 0-8.8e-5 | NA |
| % migrants | Eja | 6.91e-5 | 9.45e-5 | 8.61e-5 | NA | 5.4e-5-8.4e-5 | 7.3e-5-1.2e-4 | 3.9e-5-1.4e-4 | NA |
| % migrants | Fus | 4.94e-5 | 4.34e-5 | 1.14e-4 | NA | 4.1e-5-6.0e-5 | 3.5e-5-5.2e-5 | 9.3e-5-1.4e-4 | NA |

Divergence between the ancestral riverine and Lake Ejagham lineages was estimated to have occurred around 9.76 ka ago (95% HPD: 8.27-11.23, Fig 4A), which we consider an estimate of the timing of the colonization of Lake Ejagham. Encouragingly, this coincides with the age of the lake estimated from core samples (9 kya: Stager et al. 2017). In contrast to rapid colonization of the new lake, we estimated that the first speciation event in Lake Ejagham lineage only occurred 1.20 [0.81-1.62] ka ago, rapidly followed by the second 0.69 [0.29-1.10] ka ago. These divergence dates remained relatively similar even in models with no gene flow (point estimates 8.80, 2.15, and 1.05 ka ago, Fig 4B).

Inferred effective population sizes among Ejagham *Coptodon* varied about fourfold. We inferred a smaller effective population size for *C. ejagham* (N_e_ = 933 [406-1,524]) compared to the other two crater lake species (*C. deckerti:* 3,680 [1,249-6,539], *C. fusiforme:* 2,864 [1,514-4,743], Table 1, Fig 4E-F), which is in line with field observations of its rarity (Martin, 2013) and with its piscivorous ecology (Dunz and Schliewen, 2010).

In agreement with the results from genome-wide admixture statistics, we infer that secondary gene flow from riverine species has taken place mostly or only from *C. sp. Mamfé* relative to *C. guineensis.* In models with single migration bands, significant gene flow was inferred from *C. sp. Mamfé* into all Ejagham lineages (Fig 4D-F). Rates of gene flow to ancestral populations dropped relative to extant lineages in models with all migration bands, in particular for gene flow to the lineage ancestral to all three species (Fig 4D-F).

Overall, G-PhoCS inferred similar rates of gene flow from *C. sp. Mamfé* to extant species (Fig 4D-F). Nevertheless, due to a higher inferred rate to the *C. deckerti* - *C. ejagham* ancestor than to *C. fusiforme*, we infer that since its divergence, *C. fusiforme* experienced less gene flow than *C. deckerti* and than *C. ejagham* (40.6% and 43.2% less, respectively, in terms of the “total migration rate” estimated in single migration band models), which agrees with the result from D-statistics (Fig 3A). However, due to the higher rate inferred in the band between *C. sp. Mamfé* and the Ejagham ancestor, and the longer time span of this band, the estimated total migration rate since the split of the ancestral Ejagham lineage differs only by 6.63% between *C. fusiforme* and *C. ejagham*, 6.39% between *C. fusiforme* and *C. deckerti*, and 0.67% between *C. deckerti* and *C. ejagham* (Table 1, Fig 4D-F).

We did not find clear evidence for gene flow into Ejagham *Coptodon* from other sources besides *C. sp. Mamfé* using G-PhoCS. All rates of gene flow into Lake Ejagham lineages from *C. guineensis* or from the riverine ancestor (prior to the split between *C. sp. Mamfé* and *C. guineensis*) had 95% HPD intervals that overlapped with zero, and all except two had means very close to zero (Table 1, S4A Fig). Only the estimates of gene flow from *C. guineensis* into the two ancestral Ejagham lineages had mean population migration rates above 0.01 (0.18 and 0.47) and high variance (S4A Fig), suggesting either the possibility of low levels of ancestral gene flow from *C. guineensis*, or that gene flow from *C. guineensis* at that period may be conflated with gene flow from *C. sp. Mamfé.* In support of the latter idea, in models that combined gene flow to ancestral Ejagham lineages from *C. sp. Mamfé* and *C. guineensis*, gene flow from *C. guineensis* was again not different from zero, while variance was much smaller, and gene flow from *C. sp. Mamfé* remained significant (S4B Fig).

We also did not find clear evidence for gene flow among Ejagham *Coptodon* lineages using G-PhoCS. We evaluated models with each one of all possible migration bands in both directions, and 95% HPD for all migration rates overlapped with zero (S4C Fig). The mean inferred population migration rate was higher than 0.01 only for *C. fusiforme* to *C. deckerti* (0.27) and to *C. ejagham* (0.02). Such limited evidence for post-divergence gene flow within the radiation is surprising, given that these species are in the earliest stages of speciation (Martin, 2013). However, caution is warranted given that the very recent divergence of these lineages may render it difficult to identify ongoing gene flow. Furthermore, representative breeding pairs at the tail ends of the phenotype distribution were selectively chosen for sequencing (Martin, 2012), while excluding ambiguous individuals that could not be assigned to a particular species.

### Admixture blocks support ongoing gene flow from C. sp. Mamfé

To identify genomic blocks of admixture between riverine and Lake Ejagham species, we first defined putative blocks as contiguous sliding windows that were outliers for *f_d_*, a four-population introgression statistic related to *D* that is suitable for application to small genomic regions, and subsequently applied HybridCheck (Ward and van Oosterhout, 2016) to validate and age these blocks. We used all combinations of ingroup triplets that could differentiate between admixture from *C. guineensis* and *C. sp. Mamfé*, as well as those that could identify differential admixture among Lake Ejagham species (from either riverine species) (S8 Table). Of 1,138 putative blocks identified as *f_d_* outliers, 340 were validated by HybridCheck (93 from *C. guineensis*, and 247 from *C. sp. Mamfé*). While such blocks represent areas with ancestry patterns consistent with admixture, these patterns can also be produced by incomplete lineage sorting (ILS). To distinguish between ILS and admixture, we took advantage of our estimates of block age (coalescence time between the focal species pair) from HybridCheck and our estimates of divergence times from G-PhoCS. While nearly a quarter of blocks were estimated to be older than the Lake Ejagham lineage, and therefore likely represent ILS (S5 Fig), we identified 259 “likely” candidate regions (with a point estimate of age younger than that of the Lake Ejagham lineage), including a subset of 146 “high-confidence” regions (with non-overlapping confidence intervals of age estimates), resulting from secondary gene flow into Ejagham. In total, high-confidence admixture blocks spanned across only 0.64% of the queried part of the genome (5.7 Mb).

In accordance with the much stronger evidence for Lake Ejagham admixture with *C. sp. Mamfé* than with *C. guineensis*, the majority of likely (68.3%) and high-confidence (80.1%) admixture blocks involved *C. sp. Mamfé* as the riverine species, and likely and high-confidence admixture blocks with *C. sp. Mamfé* were, on average, younger (2.94 and 1.37 ka, respectively) than those with *C. guineensis* (4.55 and 1.97 ka, respectively, Fig 5A).

**Fig 5.**
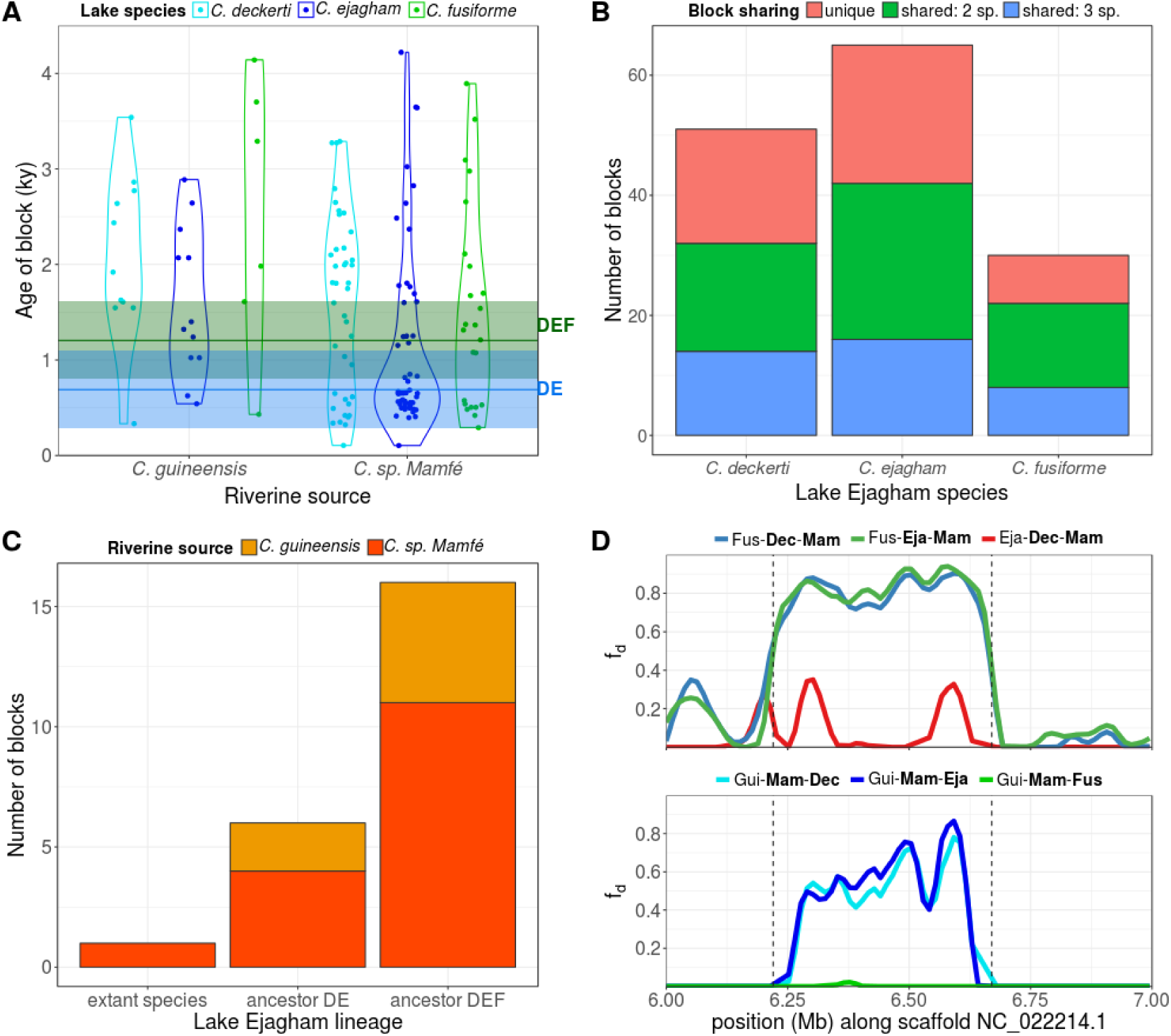
Evidence for introgression from admixture blocks. Only “high-confidence” admixture blocks, that is with a maximum estimated age younger than minimum estimated divergence time of Ejagham *Coptodon* are shown. **(A)** Age estimates of admixture blocks show ongoing introgression. Estimated divergence times of *C. deckerti* and *C. ejagham* (blue line DE), and of *C. fusiforme* and the DE ancestor (green line DEF), and the corresponding 95% HPD intervals, are also shown. **(B)** Both unique and shared (either among two or three species) admixture blocks are detected, and fewest blocks are detected in *C. fusiforme*. **(C)** A subset of blocks could be categorized using DFOIL statistics, the large majority of which introgressed to the ancestral Ejagham lineage (“ancestor DEF”). **(D)** An example of an admixture block, which is shared between *C. deckerti* and *C. ejagham*, and estimated by HybridCheck to have been introgressed 2,486 (1,651-3,554) years ago.

Because *f_d_* and HybridCheck detect admixture only between species pairs, we took two approaches to investigate at which point along the Lake Ejagham phylogeny admixture took place for likely admixture blocks. First, we intersected admixture blocks involving different Lake Ejagham species but the same riverine species, and detected 76 likely (and 38 high-confidence) blocks involving a single Lake Ejagham species, 88 (50) blocks shared among two Lake Ejagham species, and 95 (87) blocks shared among all three Lake Ejagham species (Fig 5B). Thus, 29.3% of likely blocks (and 26.0% of high-confidence blocks) were unique to a single lake species, but this may be an overestimate, since such blocks may have been present but escaped statistical detection in other species, for instance due to recombination within the block. This possibility is underscored by the age distribution of admixture blocks: admixture blocks detected in one species were not younger than those detected in multiple species (S5 Fig). In line with results from genome-wide admixture statistics and G-PhoCS, we found more admixture blocks into *C. deckerti, C. ejagham*, and their ancestor, compared to *C. fusiforme* (Fig 5B).

Second, we used D_FOIL_ statistics to distinguish between admixture involving the ancestral Lake Ejagham lineage (“DEF”), the *C. deckerti* – *C. ejagham* ancestor (“DE”), and the terminal branches. We were able to categorize 23 likely (and 13 high-confidence) admixture blocks with D_foil_ statistics, showing a pattern of decreasing occurrence of admixture blocks through time, with only a single likely (and 0 high-confidence) block involving a terminal Lake Ejagham branch (Fig 5C). For cases where admixture is with an ancestral (lake) clade, D_FOIL_ statistics cannot infer the direction of introgression, but the single classified admixture block with an extant lake taxon is, as expected, inferred to have been into the lake.

### Admixture of olfactory genes into C. deckerti and C. ejagham

Among all high-confidence blocks, 11 gene ontology terms were enriched (Table 2). Eight genes in a single admixture block on scaffold NC_022214.1 were responsible for the three most enriched categories; seven of these genes are characterized as olfactory receptors and the eighth as “olfactory receptor-like protein” (none have a gene name and only one has 1-to-1 orthologues in other species on Ensembl Release 90 (S10 Table)). The admixture block containing this cluster of genes, which is shown in Fig 5D, was estimated to have introgressed from *C. sp. Mamfé* into both *C. deckerti* and *C. ejagham* 2,486 (1,651-3,554) years ago, shortly prior to the divergence of the *C. deckerti* / *C. ejagham* ancestor from *C. fusiforme*, 1,205 (806-1,616) years ago. Among all high-confidence admixture blocks, this block was the largest, had the highest summed *f_d_* score, and had the second lowest HybridCheck p-value.

**Table 2.**
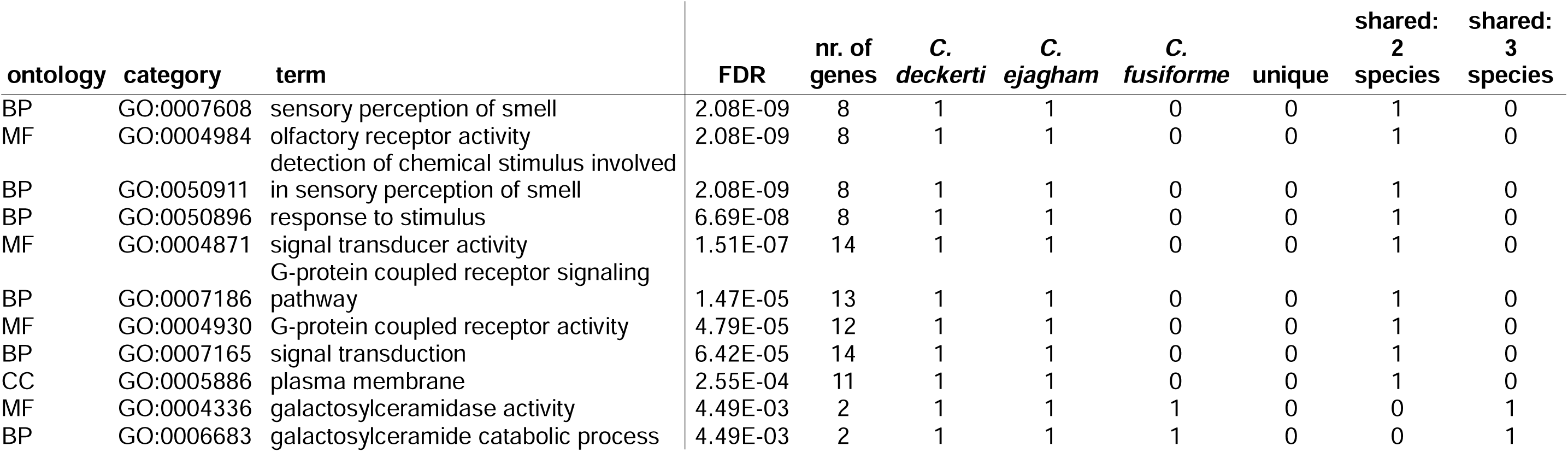
**Gene Ontology term enrichment among genes in admixture blocks.** FDR and number of genes are given for genes in all “high-confidence” admixture blocks. The last six columns indicate whether (1) or not (0) each term was also enriched (FDR < 0.05) for subsets of admixture blocks involving each species and each block sharing category (“unique” – blocks unique to one Lake Ejagham species; “shared: 2/3 species” – blocks shared among two/three Lake Ejagham species. No additional GO terms were enriched for admixture blocks subsets only. Ontologies: BP = Biological Process, CC = Cellular Component, MF = Molecular Function.

When performing GO analyses separately for blocks involving each Lake Ejagham species, no additional terms were found to be enriched. With respect to admixture blocks involving each Lake Ejagham species, the same 11 terms were enriched for *C. ejagham*, nine of these terms were enriched for *C. deckerti*, and none were enriched for *C. fusiforme* (Table 2). Blocks unique to one Lake Ejagham species (either taken together, or separately by species), were not enriched for any terms, while blocks shared between two species were enriched for nine terms and blocks shared between all three species for two terms (Table 2).

## Discussion

In the context of an isolated lake, a classic case of fully sympatric speciation would involve i) colonization of the lake by a single lineage, effectively in a single event, and ii) no subsequent gene flow with populations outside of the lake prior to or during speciation. Our results suggest that for the Lake Ejagham *Coptodon* radiation, the former is true but the latter is not. In contrast to the original paradigm of a highly isolated lake colonized only once by a single cichlid pair (Schliewen et al. 2001), we found ongoing gene flow from one of the riverine species into all three species in the lake throughout their speciation histories. Interestingly, one of the clearest signals of introgression came from a cluster of olfactory receptor genes that introgressed into the ancestral population at xx kya just prior to the first speciation event, suggesting that gene flow may have facilitated speciation.

### Rapid initial colonization of Lake Ejagham

Our estimates of the origin of the Ejagham Coptodon lineage (9.76 ka ago, Fig. 4A) were nearly identical to the estimated date of the origin of the lake itself (9 ka years ago, Stager et al., 2017), suggesting that the lake was rapidly colonized by the ancestral lineage. It should be noted that this estimate in turn relies on an estimate of the mutation rate. We here use estimates from sticklebacks (Guo et al., 2013) as previous studies on cichlids have done (Kautt et al., 2016a, 2016b), but it cannot be excluded that our focal species have substantially different mutation rates (Martin and Höhna, 2017; Martin et al., 2017; Recknagel et al., 2013).

Martin et al. (2015a) argued that the Cameroon lakes containing cichlid radiations may not be as isolated as has previously been suggested, based on the inference of secondary gene flow in all radiations, and the fact that each lake has been colonized by several fish lineages (five in the case of Lake Ejagham). Our inference of a rapid, successful colonization process and evidence for ongoing gene flow are both in support of this view. In this light, it is worth pointing out that lake Ejagham (i) has an outflow in the wet season (S6 Fig) which may be connected to the Munaya River (itself part of the Cross River system), (ii) does *not* have a waterfall that could prevent fish from entering the lake (C. H. Martin, pers. obs.) contrary to claims elsewhere (Bolnick and Fitzpatrick, 2007), and (iii) is at an elevation of only 141 meters, about 60 meters higher than the closest river drainage (the other two Cameroon lakes containing sympatric radiations, Lake Barombi Mbo and Lake Beme, are at altitudes of 314 and 472 meters, respectively).

### No major secondary colonizations

Our data suggest that the initial colonization of the lake established the large majority of the lineage that has since diversified within Lake Ejagham, and we find no evidence for major secondary colonizations that either established a new lineage or resulted in a hybrid swarm. Several lines of evidence indicate that such events are unlikely to have taken place. First, considerable phylogenetic conflict would be expected if diversification happened rapidly after a secondary colonization event, while we found particularly widespread monophyly across the genome (89.34%, S7 Table). Second, we inferred a long time lag between colonization and the first speciation event within the lake (9.76 ka and 1.20 ka ago, respectively, Fig 4A, Table 1). Third, we estimated gene flow into the ancestral lake lineage to be relatively low (Fig 4B), and in line with this, models with and without post-divergence gene flow between riverine and lake lineages resulted in similar (9.76 and 8.80 ka ago, respectively, Table 1) estimates of the divergence time of the ancestral lake lineage.

### Continuous low levels of gene flow from one of two Cross River Coptodon species

Even though we found that Ejagham *Coptodon* was established by a single major colonization, our results are not consistent with subsequent isolation of the lake lineages. We found evidence for ongoing secondary gene flow from the source population, which diverged into *C. guineensis* and *C. sp. Mamfé* after the split with the Ejagham lineage. Results from all three types of approaches that we used to identify secondary gene flow (demographic analysis with G-PhoCS, genome-wide admixture statistics, and the identification of admixture blocks) show that gene flow originated predominantly from one of these riverine lineages, *C. sp. Mamfé* (Fig 3A, 4B, 5). Little is known about the precise geographic distribution of *C. sp. Mamfé*, yet this asymmetry is consistent with the sampling location of this species (37 km from Lake Ejagham to the Cross River at Mamfe) relative to that of *C. guineensis* (65 km from Lake Ejagham to a tributary of the Cross River at Nguti; see also Fig 1 that depicts all major rivers). Both *Coptodon* lineages are known to coexist within the Cross River drainage. Our data suggest that C. sp. Mamfe is most likely a new species.

Evidence for gene flow from *C. guineensis* was much weaker compared to *C. sp. Mamfé* and was mostly restricted to ancestral Lake Ejagham lineages (admixture blocks: Fig 5, G-PhoCS: S4A-B Fig). It should furthermore be noted that the assignment of the riverine source lineage is likely to be more error-prone further back in time, given the recent divergence between *C. guineensis* and *C. sp. Mamfé*. However, the clearest evidence of gene flow from *C. guineensis* comes from admixture blocks, where an inference of differential ancestry from the two riverine species was required. Since we were only able to include a single *C. sp. Mamfé* individual, it is nevertheless possible that we missed genetic variation in that species, which may have led to incorrect assignment as the riverine source lineage as *C. guineensis*.

### Differential admixture of Ejagham radiation with riverine Coptodon

We found some evidence for differential riverine admixture, from *C. sp. Mamfé*, among the three Ejagham species. While the admixture proportion of *C. ejagham* may be slightly higher than that of *C. deckerti* (*f_4_*-ratio test: Fig 3B, but see D-statistics, Fig 3A, and G-PhoCS: Fig 4A-C), the evidence was stronger for elevated riverine admixture with sister species *C. deckerti* and *C. ejagham* relative to *C. fusiforme* (D-statistics: Fig 3A, admixture blocks: Fig 5), which specifically appears to originate from high admixture with the *C. deckerti* / *C. ejagham* ancestor (G-PhoCS: Fig 4B). In accordance with this, Martin et al. (2015a) identified riverine admixture with the *C. deckerti*/*C. ejagham* ancestor using Treemix. Martin et al. (2015a) found that a proportion of *C. fusiforme* individuals appeared more admixed than any other Ejagham *Coptodon.* The magnitude of the effect in their PCA plot (Fig 3C in Martin et al., 2015a), as well as the fact that only some of the *C. fusiforme* individuals were involved, suggests contemporary hybridization; however, this was not supported by their STRUCTURE analysis of the same data. Nonetheless, contemporary hybridization may have resulted from the known introduction of riverine fishes into this lake by a local town council member in 2000-2001 (Martin, 2012). This resulted in the establishment of a *Parauchenoglanis* catfish within the lake, still recorded as abundant in 2016 (CHM pers. obs.). However, no riverine *Coptodon* have been confirmed beyond a posted sign reporting introduced river fishes. In this study, we found no evidence that any of our individuals were recent hybrids (Fig S4), but our limited sample sizes preclude us from any strong inferences on their potential occurrence in the lake.

### Introgression of a cluster of olfactory receptor genes shortly prior to speciation

Complex patterns of secondary gene flow such as those observed here are not easily interpreted in terms of their contribution to speciation. The formation of hybrid swarms has been suggested to promote speciation (Kautt et al., 2016a; Seehausen, 2004), yet we did not find evidence for major secondary colonizations, or a specific admixture event that could be linked to the timing of speciation. Instead, we inferred ongoing gene flow, which could theoretically inhibit speciation, by counteracting incipient divergence within the lake, or promote speciation, by introducing novel genetic variation or co-adapted gene complexes.

Interestingly, one admixture block contained a cluster of eight olfactory receptor genes (S10 Table), causing a highly significant overrepresentation of several gene ontology terms containing these genes (Table 2). While in mammals, the Olfactory Receptor (OR) gene family is the largest gene family with around 1,000 genes, mostly due to the expansion of a single group of genes, fish species examined so far have much fewer (69-158 complete genes) yet a more diverse set of OR genes (Azzouzi et al., 2014; Niimura and Nei, 2005). Unfortunately, little additional information is known about the eight admixed olfactory receptor genes.

This cluster of OR genes was contained in the largest and arguably most striking of all high-confidence admixture blocks (Fig 5D), which is estimated to have introgressed from *C. sp. Mamfé* into *C. deckerti* and *C. ejagham*, but not *C. fusiforme*, just prior to the estimated divergence time of *C. fusiforme* and the ancestor of *C. deckerti* and *C. ejagham.* Thus, the timing, source and target of introgression all correspond with the inference of elevated levels of gene flow from *C. sp. Mamfé* to the *C. deckerti - C. ejagham* ancestor relative to *C. fusiforme* (Fig 3A, Fig 4B, Fig 5). These patterns may suggest a role for the introgression of this block in initiating speciation in Ejagham *Coptodon*.

Chemosensory signaling in general, and olfactory receptors specifically, have often been linked to speciation, especially with respect to sexual isolation (Smadja and Butlin, 2008). A host of studies has shown the importance of olfactory signaling in conspecific mate recognition in fish (Crapon de Caprona and Ryan, 1990; Kodric-Brown and Strecker, 2001; McLennan, 2004; McLennan and Ryan, 1999), and in a pair of closely related Lake Malawi cichlids, female preference for conspecific males was shown to rely predominantly if not exclusively on olfactory cues (Plenderleith et al., 2005). Not surprisingly, it has repeatedly been suggested that olfactory signals may help explain explosive speciation in cichlids (Azzouzi et al., 2014; Blais et al., 2009; Keller-Costa et al., 2015).

Olfactory signaling seems particularly relevant to mate choice and speciation in Ejagham *Coptodon*, since three species occur syntopically, assortative mating by species appears to represent the strongest isolating barrier (Martin, 2012, 2013), and sexual dichromatism is absent. Important next steps will be to examine the importance of olfactory cues in mate recognition in Lake Ejagham *Coptodon*, specifically between *C. fusiforme* and the other two species, and to characterize these genes and their patterns of divergence and admixture in more detail.

### Waiting time for sympatric speciation

While we inferred that colonization of Lake Ejagham took place more than 9 ka years ago, the first branching event among Ejagham *Coptodon* was estimated to have occurred as recently as 1.20 ka years ago (Fig 4A, Table 1). We did not include the fourth nominal *Coptodon* species in the lake, *C. nigrans*, but extreme phenotypic similarity to *C. deckerti* (Dunz and Schliewen, 2010) and our inability to identify or distinguish these individuals in field collections and observations (Martin 2012, 2013) suggests a close relationship between *C. deckerti* and this nominal species, which would not change this inference. It thus appears that during the large majority of the time that the *Coptodon* lineage was present in Lake Ejagham, no diversification occurred. One possibility is that earlier speciation events did occur, but were followed by extinction. While we cannot fully exclude this scenario, there are no indications for environmental disruptions such as major changes in water chemistry or depth during the history of Lake Ejagham (Stager et al., 2017).

Assuming that the divergence of *C. fusiforme* was the first within this radiation, a striking difference emerges between the waiting time to the first (7.74 ka) and the next two speciation events, which both occurred within 1.20 ka. The opposite pattern, a slowing speciation rate, would be expected if speciation followed a niche-filling model of ecological opportunity in the lake. At least two non-mutually exclusive explanations may account for this counterintuitive result.

First, an initial lack of ecological opportunity in young Lake Ejagham may have prevented a rapid first speciation event. Our results are reminiscent of those for sympatrically speciating Tristan da Cunha buntings (Ryan et al., 2007), where, as discussed by Grant and Grant (2009), the ancestral branch is considerably longer than those of the extant species. Grant and Grant (2009) propose that plants that constitute one of the niches used by the extant finch species may have arrived only recently. Similarly, ecological diversity in lower trophic levels in the lake may have been insufficient to generate the necessary degree of disruptive selection to drive divergence. For instance, *Daphnia* never colonized another Cameroon crater lake, Barombi Mbo, during its ca. 1 million year existence (Cornen et al., 1992; Green and Kling, 1988).

Second, genetic variation for traits underlying sexual and ecological selection and the associated genetic architecture may initially not have been conducive to speciation. If ecological and mate preference traits are distinct (i.e. not magic traits: Servedio et al., 2011) and independently segregating within the ancestral colonizing population, sympatric speciation models predict that there will be a waiting time associated with the initial buildup of linkage disequilibrium between these traits before sympatric divergence can proceed (Dieckmann and Doebeli, 1999; Kondrashov and Kondrashov, 1999). Furthermore, Bolnick (2004) demonstrated that under conditions where genetic variation for stringent assortative mating is limiting and females are penalized for assortative mating, sympatric speciation may require a long time. In this light, it is particularly intriguing that introgression of a block containing eight olfactory receptor genes from *C. sp. Mamfé*, which are likely to be highly relevant for mate choice, were introgressed shortly prior to the first speciation event. Therefore, genetic variation brought in by riverine gene flow may have been necessary to initiate speciation among Lake Ejagham *Coptodon*.

### Conclusions

We showed that Lake Ejagham was rapidly colonized by ancestors of the extant *Coptodon* radiation in a single major colonization, while also inferring low levels of ongoing and continuous secondary gene flow from riverine species into ancestral as well as extant lake species. Speciation can still be considered sympatric if secondary gene flow was present but did not play a causal role in speciation, and the pattern of ongoing gene flow is consistent with this. However, introgression of a cluster of olfactory receptor genes into a pair of sister species (but not the third species) just prior to their divergence, indicates that secondary gene flow may have been important to speciation. The introgression of olfactory genes is particularly salient given that Ejagham *Coptodon* species exhibit strong assortative mating, but currently weak disruptive selection, syntopic breeding territories, and no sexual dichromatism within a tiny, shallow lake.

## Methods

### Sampling

Sampling efforts and procedures have been described previously in Martin et al. (2015a). Here, we sampled breeding individuals displaying reproductive coloration from three species of *Coptodon* (formerly *Tilapia*) that are endemic to Lake Ejagham in Cameroon: *Coptodon fusiforme* (n = 3), *C. deckerti* (n = 2), and *C. ejagham* (n = 2). We additionally used samples from closely related riverine species from the nearby Cross River whose ancestors likely colonized Lake Ejagham: *C. guineensis* (n = 2) at Nguti, 65 km from Lake Ejagham, and an undescribed taxon, *C. sp. “Mamfé”* (Keijman, 2010) (n = 1), at Mamfé, 37 km from Lake Ejagham. Finally, we sampled a closely related outgroup species, *C. kottae*, from crater lake Barombi ba Kotto (145 km from Lake Ejagham), and a distantly related outgroup species, *Sarotherodon galilaeus* (n = 3), from the Cross River at Mamfé. Cichlids were caught by seine or cast-net in 2010 and euthanized in an overdose of buffered MS-222 (Finquel, Inc.) following approved protocols from University of California, Davis Institutional Animal Care and Use Committee (#17455) and University of North Carolina Animal Care and Use Committee (#15-179.0), and stored in 95-100% ethanol or RNAlater in the field.

### Genome sequencing and variant calling

DNA was extracted from muscle tissue using DNeasy Blood and Tissue kits (Qiagen, Inc.) and quantified on a Qubit 3.0 fluorometer (Thermofisher Scientific, Inc.). Genomic libraries were prepared using the automated Apollo 324 system (WaterGen Biosystems, Inc.) at the Vincent J. Coates Genomic Sequencing Center (QB3). Samples were fragmented using Covaris sonication, barcoded with Illumina indices, and quality checked using a Fragment Analyzer (Advanced Analytical Technologies, Inc.). Nine to twelve samples were pooled in four different libraries for 150PE sequencing on four lanes of an Illumina Hiseq4000.

We mapped raw sequencing reads in fastq format to the *Oreochromis niloticus* genome assembly (version 1.1, https://www.ncbi.nlm.nih.gov/assembly/GCF_000188235.2/, Brawand et al., 2014) with BWA-MEM (version 0.7.15, Li, 2013). Using Picard Tools (version 2.10.3, http://broadinstitute.github.io/picard), the resulting .sam files were sorted (*SortSam* tool), and the resulting .bam files were marked for duplicate reads (*MarkDuplicates* tool) and indexed (*BuildBamlndex* tool). SNPs were called using the HaplotypeCaller program in the Genome Analysis Toolkit (GATK; DePristo et al., 2011), following the GATK Best Practices guidelines (Van der Auwera et al., 2013), https://software.broadinstitute.org/gatk/best-practices/). Since no high-quality known variants are available to recalibrate base quality and variant scores, SNPs were called using hard filtering in accordance with the GATK guidelines (DePristo et al., 2011; Van der Auwera et al., 2013): QD < 2.0, MQ < 40.0, FS > 60.0, SOR > 3.0, MQRankSum < -12.5, ReadPosRankSum < -8.0. SNPs that did not pass these filters were removed from the resulting VCF files using vcftools (version 0.1.14, Danecek et al., 2011, using “--remove-filtered-all” flag), as were SNPs that differed from the reference but not among focal samples (using “max-non-ref-af 0.99” in vcftools) and SNPs with more than two alleles (using “-m2 -M2” flags in bcftools, version 1.5 (Li, 2011)). Genotypes with a genotype quality below 20 and depth below 5 were set to missing (using “--minGQ” and “--minDP” flags in vcftools, respectively), and sites with more than 50% missing data were removed (using “--max-missing” flag in vcftools). Our final dataset consisted of 15,523,738 SNPs with a mean sequencing depth of 11.82 (range: 7.20 – 16.83) per individual.

### Phylogenetic trees, networks, and genetic structure

We employed several approaches to estimate relationships among the three species in the *Coptodon* Ejagham radiation and the two riverine *Coptodon* taxa. These analyses were repeated for four outgroup configurations: no outgroup (unrooted trees), using only *C. kottae*, only *S. galilaeus*, and both *C. kottae* and *S. galilaeus* as outgroups. Only sites with less than 10% missing data were used for phylogenetic reconstruction.

Using the GTR-CAT maximum likelihood model without rate heterogeneity, as implemented in RaxML (version 8.2.10, Stamatakis, 2014), we inferred phylogenies for all SNPs concatenated, as well as separately for each 100kb window with at least 250 variable sites (“gene trees”). This resulted in sets of 1,532 – 2,559 trees, depending on the outgroup configuration.

Next, rooted gene trees were used, first, to compute Internode Confidence All (ICA) scores (Salichos et al., 2014), using the “-L MR” flag in RaXML) for each of the nodes of the whole-genome trees. Rooted gene trees were also used to construct species trees in Phylonet (version 3.6.1, Than et al., 2008) using the Minimize Deep Coalescence criterion (Than and Nakhleh, 2009, “Infer_ST_MDC” command) and maximum likelihood (“Infer_Network_ML” command with zero reticulations), and using a maximum pseudo-likelihood method implemented in MP-EST (version 1.5, Liu et al., 2010). Finally, we used ASTRAL (version 2.5.5, Mirarab et al., 2014) to infer species trees from unrooted gene trees.

To visualize patterns of genealogical concordance and discordance, we computed a phylogenetic network using the NeighborNet method (Bryant and Moulton, 2004) implemented in Splitstree (version 4.14.4, Huson and Bryant, 2006), using all SNPs.

We used the machine learning program *Saguaro* (Zamani et al., 2013) to determine the dominant topology across the genome and calculate the percentages of the genome that supported specific relationships, such as monophyly of the Ejagham *Coptodon* radiation. *Saguaro* combines a hidden Markov model with a self-organizing map to characterize local phylogenetic relationships among individuals without requiring a priori hypotheses about the relationships. This method infers local relationships among individuals in the form of genetic distance matrices and assigns segments across the genomes to these topologies. These genetic distance matrices can then be transformed into neighborhood joining trees to visualize patterns of evolutionary relatedness across the genome. To be comprehensive in our search, we allowed Saguaro to propose 31 topologies for the genome, but otherwise applied default parameters. We investigated the effect of the number of proposed topologies on the proportion of genomes assigned to our two categories, and found that the percentages were robust after 20 proposed topologies, with increasingly smaller percentages of the genome being assigned to new additional topologies.

### Genome-wide tests for admixture

We tested for admixture between the two riverine species and the three Lake Ejagham species using several statistics based on patterns of derived allele sharing among these species. We used the ADMIXTOOLS (version 4.1, Patterson et al., 2012) suite of programs to compute four-taxon D-statistics (“ABBA-BABA tests”, *qpDstat* program) and a five-taxon *f_4_*-ratio test (*qpF4ratio* program), and the software dfoil (release 2017-06-14, http://www.github.com/jbpease/dfoil, Pease and Hahn, 2015) to compute five-taxon D_FOIL_ statistics. For all analyses, we used *S. galilaeus* as the outgroup species.

Given a topology (((P1, P2), P3), O), *D* can identify admixture between either P1 or P2 on one hand, and P3 on the other based on the relative occurrence of ABBA and BABA patterns. First, we computed D-statistics to test for admixture between *C. guineensis* (P1) or *C. sp. Mamfé* (P2) and any Lake Ejagham species (P3). Given that all three of these comparisons indicated admixture between *C. sp. Mamfé* and Lake Ejagham species (Fig 3A), we next tested whether there was evidence for differential admixture from *C. sp. Mamfé* among the three Ejagham *Coptodon* species, using the three possible pairs of Lake Ejagham species as P1 and P2, and *C. sp. Mamfé as P3*.

Another way to test for differential *C. sp. Mamfé* admixture among Ejagham *Coptodon* species is by using *f_4_*-ratio tests, wherein taxon “X” is considered putatively admixed, containing ancestry proportion α from the branch leading to P2 (after its divergence from taxon P1), and ancestry proportion α – 1 from the branch leading to taxon P3. Given the constraints imposed by the topology of our phylogeny, we could only test for admixed ancestry of either *C. deckerti* or *C. ejagham* with *C. sp. Mamfé*, after divergence of the *C. deckerti – C. ejagham* ancestor from *C. fusiforme*. Testing for admixed ancestry of *C. fusiforme* using an *f_4_*-ratio test would merely produce a lower bound of α (see Mailund, 2014), while we were instead interested in an estimate or upper bound on α, since our null hypothesis was α = 1: *C. fusiforme* has ancestry only from the *C. deckerti – C. ejagham* ancestor.

The five-taxon D_FOIL_ statistics enable testing of the timing, and in some cases, direction of introgression in a symmetric phylogeny with two pairs of taxa with a sister relationship within the provided phylogeny, and an outgroup. Given our six-taxon phylogeny, we performed this test for three sets of five species, each with a unique combination of two of the three Ejagham *Coptodon* species as one species pair (P1 and P2), and *C. guineensis* and *C. sp. Mamfé* as the second species pair (P3 and P4; the outgroup again being *S. galilaeus*). The test involves the computation of four D_FOIL_ statistics (D_FO_, D_IL_, D_FI_, and D_OL_), each essentially performing a three-taxon comparison. The combination of results for these statistics can inform whether introgression predominantly occurred among any of the four ingroup extant taxa, in which case the direction of introgression can also be inferred (e.g. P1 → P3), or among an extant taxon and the ancestor of the other species pair, in which case the direction of introgression cannot be inferred (e.g. P1 ↔ P3P4). Unlike D and *f_d_* statistics, D_FOIL_ statistics by default also include counts of patterns where only a single taxon has the derived allele (e.g. BAAAA), under the assumption of similar branch lengths across taxa. When this assumption is violated, the dfoil program can be run in “dfoilalt” mode, thereby excluding single derived-allele counts (Pease and Hahn, 2015). Since we observed significantly fewer single derived-allele sites for *C. sp. Mamfé* than for *C. guineensis*, we ran the dfoil program in “dfoilalt” mode at a significance level of 0.001.

### Inference of demographic history with G-PhoCS

For a detailed reconstruction of the demographic history of Ejagham *Coptodon* and the two closely related riverine species, we used the program G-PhoCS (Generalized Phylogenetic Coalescent Sampler, version 1.3, Gronau et al., 2011). G-PhoCS implements a coalescent-based approach using Markov chain Monte Carlo (MCMC) to jointly infer population sizes, divergence times, and optionally migration rates among extant as well as ancestral populations, given a predefined population phylogeny. To infer migration rates, one or more unidirectional migration bands can be added to the model, each between a pair of populations that overlap in time. G-PhoCS can thus infer the timing of migration within the bounds presented by the population splits in the phylogeny.

As input, G-PhoCS expects full sequence data for any number of loci. Since G-PhoCS models the coalescent process without incorporating recombination, it assumes no recombination within loci, and free recombination between loci. Following several other studies (Choi et al., 2017; Gronau et al., 2011; Hung et al., 2014; McManus et al., 2015), we picked 1 kb loci separated by at least 50 kb. Following (Gronau et al., 2011), loci were selected not to contain the following classes of sites within the *O. niloticus* reference genome – that is, rather than being simply masked, these sites were not allowed to occur in input loci: (1) hard-masked (N) or soft-masked (lowercase bases) sites in the publicly available genome assembly; (2) sites that were identified to be prone to ambiguous read mapping using the program SNPable (Li, 2009, using k=50 and r=0.5 and excluding rankings 0 and 1); and (3) any site within an exon or less than 500bp from an exon boundary. Furthermore, loci were chosen to contain no more than 25% missing data (uncalled and masked genotypes). Using these selection procedures, a total of 2,618 loci were chosen using custom scripts and a VCF to Fasta conversion tool (Bergey, 2012).

Prior distributions for demographic parameters are specified in G-PhoCS using α and β parameters of a gamma distribution. We determined the mean of the prior distribution (α / β) for each parameter using a number of preliminary runs, while keeping the variance (α / β^2^) large following (Gronau et al., 2011) to minimize the impact of the prior on the posterior (see S9 Table for all G-PhoCS settings). Preliminary runs confirmed that regardless of the choice of the prior mean, MCMC runs converged on similar posterior distributions.

For each combination of migration bands (see below), we performed four replicate runs. Each G-PhoCS run was allowed to continue for a week on 8-12 cores on a single 2.93 GHz compute node of the UNC Killdevil computing cluster, resulting in runs with 1-1.5 million iterations. The first 250,000 iterations were discarded as burn-in, and the remaining iterations were sampled 1 in every 50 iterations. Convergence, stationarity, and mixing of MCMC chains was assessed using Tracer (version 1.6.0, Rambaut et al., 2014).

Because the total number of possible migration bands in a six-taxon phylogeny is prohibitively high for effective parameter inference, we took the following strategy. Our primary focus was on testing migration bands from *C. sp. Mamfé* (“Mam”) and *C. guineensis* (“Gui”) to the Lake Ejagham *Coptodon* species and their ancestors: *C. deckerti* (“Dec”), *C. ejagham* (“Eja”), *C. fusiforme* (“Fus”), “DE” (the ancestor to Dec and Eja), and “DEF” (the ancestor to DE and Fus). We first performed runs each with a single one of these migration bands. Since all migration bands from *C. sp. Mamfé* had non-zero migration rates, we next performed runs with all of these migration bands at once. However, in those runs we observed failures to converge, higher variance in parameter estimates, and the dropping to zero of rates of migration to the ancestral Lake Ejagham lineage (see Fig 4). The latter is surprising given that for single-band runs, this migration rate was the highest inferred, and is also in sharp contrast to other analyses that show much stronger support for migration to the ancestral lineage than to extant species. While we suspect that runs with all migration bands have poor performance due to the number of parameters, runs with single migration bands may be prone to overestimation of the migration rate. We therefore also performed runs with migration bands either to all three extant species or to both ancestral lineages, and report results for all of these run types, separately.

Finally, we performed runs with no migration bands. We did not examine models with migration from the Ejagham radiation to neighboring rivers because this is not relevant to sympatric speciation scenarios in this lake.

To convert the θ (4 * N_e_ * μ) and т (T * μ) parameters reported by G-PhoCS, which are scaled by the mutation rate, to population sizes N_e_ and divergence times T, we used a per year mutation rate μ of 7.5 * 10^−9^, based on a per-generation mutation rate of 7.5 * 10^−9^ (Guo et al., 2013) and a generation time of 1 year similar to East African cichlids and corresponding to observations of laboratory growth rates (although note that these species have never been bred in captivity). We converted the migration rate parameter *m* for a given migration band to several more readily interpretable statistics. First, the population migration rate (2*Nm*) is twice the number of migrants in the source population that arrived by migration from the target population, per generation. It is calculated using the value of θ for the target population (*2Nm*_s→t_ = *m*_s→t_ * θ_t_/4), and as such it does not depend on an estimate of the mutation rate. Second, the proportion of migrants per generation is calculated by multiplying *m* by the mutation rate. Third, the “total migration rate” *M* (Gronau et al., 2011) can be interpreted as the probability that a locus in the target population has experienced migration from the source population, and is calculated by multiplying *m* by the time span of the migration band, which is the time window during which both focal populations existed (*M*_s→t_ = *m*_s→t_ * т_s,t_).

### Local admixture tests

To identify genomic regions with evidence for admixture between one of the riverine species and one or more of the Lake Ejagham species, we first computed the *f_d_* statistic (Martin et al., 2015b) along sliding windows of 50 kb with a step size of 5 kb, using ABBABABA.py (Martin, 2015). The *f_d_* statistic is a modified version of the Green et al. (2010) estimator of the proportion of introgression (*f*), and has been shown to outperform *D* for the detection of introgression in small genomic windows (Martin et al., 2015b).

In the topology ((P1, P2), P3), O), *f_d_* tests for introgression between P2 and P3. For each window, *f_d_* was calculated for two types of configurations. First, those that can identify the source of any riverine admixture, using the two riverine species as P1 and P2 and a Lake Ejagham species as P3 (for example: P1 = *C. guineensis*, P2 = *C. sp. Mamfé*, P3 = *C. ejagham*). Second, those that can identify differential admixture from a riverine species among two Lake Ejagham species (for example: P1 = *C. deckerti*, P2 = *C. ejagham*, P3 = *C. sp. Mamfé*). Since *f_d_* only detects introgression between P2 and P3, *f_d_* was also computed for every triplet with P1 and P2 swapped.

P-values for *f_d_* were estimated by Z-transforming single-window *f_d_* values based on a standard normal distribution, followed by multiple testing correction using the false discovery rate method (FDR, Benjamni and Hochberg, 1995), using a significance level of 0.05. Next, putative admixture blocks were defined by combining runs of significant *f_d_* values that were consecutive or separated by at most three non-significant (FDR > 0.05) windows. Because any secondary admixture must have occurred within the last ~10k years, after colonization of Lake Ejagham, true admixture blocks are expected to be large, and blocks of less than five total windows or with maximum *f_d_* values below 0.5 were excluded from consideration. Therefore, only genomic scaffolds of at least 70 kb (i.e., 557 scaffolds or 97.40% of the assembled genome) can harbor a putative admixture block. Blocks indicating differential admixture with a riverine species among two Lake Ejagham species (in ingroup triplets with a pair of Lake Ejagham species as P1 and P2, and a riverine species as P3) were retained only when the riverine source of admixture could be distinguished in a direct comparison, by intersection with blocks indicating differential admixture with a Lake Ejagham species among the two riverine species. For instance, a block indicating admixture between *C. deckerti* (P2) and *C. sp. Mamfé* (P3) in an ingroup triplet with *C. ejagham* as P1 (i.e., identifying differential admixture among two lake species) was only retained if it overlapped with an admixture block with *C. guineensis* as P1, *C. sp. Mamfé* as P2, and *C. deckerti* as P3 (i.e. identifying differential admixture among the riverine sources with the same lake species). Putative admixture blocks as defined by *f_d_* values were validated and aged using HybridCheck (Ward and van Oosterhout, 2016), using the same mutation rate as for our G-PhoCS analysis. HybridCheck identifies blocks that may have admixed between two sequences by comparing sequence similarity between triplets of individuals along sliding windows, and next estimates, for each block, the coalescent time between the two potentially admixed sequences. While HybridCheck can also discover admixture blocks *ab initio*, we employed it to test user defined blocks with the “addUserBlock” method. Given that HybridCheck accepts triplets of individuals, and *f_d_* blocks detected in a given species triplet were tested twice in HybridCheck for that species triplet, each using a different individual of the admixed Lake Ejagham species. Blocks were retained when HybridCheck reported admixture between the same pair of individuals as the *f_d_* statistic, and with a p-value smaller than 0.001 for both triplets of individuals. Our final set of “likely blocks” consisted of those with an estimated age smaller than the G-PhoCS point estimate (in runs with all possible migration bands from *C. sp. Mamfé*) of the divergence time between the Lake Ejagham ancestor (“DEF”) and the riverine ancestor (“AU”), while “high confidence blocks” were defined as those with the *upper* bound of the 95% confidence interval of the age estimate smaller than the *lower bound* of the 95% HPD of the divergence time estimate between DEF and AU (for whichever set of G-PhoCS runs, either with no, some, or all migration bands from *C. sp. Mamfé*, had the lowest value for this parameter).

In order to characterize the patterns of admixture for these pairwise admixture blocks further, we calculated localized D_FOIL_ statistics for each. Since these statistics depend on the occurrence of sufficient numbers of all possible four-taxon derived allele frequency occurrence patterns among five taxa, these only produced results for a subset of blocks (for the same reason, we were not able to use these statistics for *ab initio* admixture block discovery along sliding windows). Since we already established the presence of admixture for these blocks, and performed these analyses to determine the pattern of admixture, we did not require significance for each D_FOIL_ statistic, but also considered it to be positive or negative if the statistic was more than half its maximum value and had at least 10 informative sites.

### Gene Ontology for admixture blocks

We assessed whether “high confidence” admixture blocks were enriched for specific gene categories using Gene Ontology (GO) analyses. Entrez Gene gene identifiers were extracted by intersecting the genomic coordinates of admixture blocks with a GFF file containing the genome annotation for *O. niloticus* (Annotation Release 102, available at https://www.ncbi.nlm.nih.gov/genome/annotationeuk/Oreochromisniloticus/102/), and GO annotations for each gene were collected using the R/Bioconductor package biomaRt (Durinck et al., 2009). Next, GO enrichment analysis was carried out with the R/Bioconductor package goseq (Young et al., 2010), using a flat probability weighting function, the Wallenius method for calculating enrichment scores, and correcting p-values for multiple testing using the false discovery rate method (FDR, Benjamni and Hochberg, 1995). GO terms were considered enriched for FDRs below 0.005.

## Acknowledgements

This study was funded by a National Geographic Society Young Explorer’s Grant, a Lewis and Clark Field Research grant from the American Philosophical Society, and the University of North Carolina at Chapel Hill to CHM. We gratefully acknowledge the Cameroonian government and the regional authority and village council of Eyumojock and surrounding communities for permission to conduct this research. We thank Cyrille Dening Touokong, Jackson Waite-Himmelwright, and Patrick Enyang for field assistance and Nono LeGrand Gonwuou for help obtaining permits.

## Data availability statement

All sequencing data will be deposited in NCBI’s Short Read Archive. Scripts and analyses output will be deposited in the Dryad Digital Repository.

## Funding

This study was funded by a National Geographic Society Young Explorer’s Grant, a Lewis and Clark Field Research grant from the American Philosophical Society, and the University of North Carolina at Chapel Hill to CHM.

## Competing interests

The authors declare no competing interests.

## Supplementary Figures & Tables

**S1 Fig.**
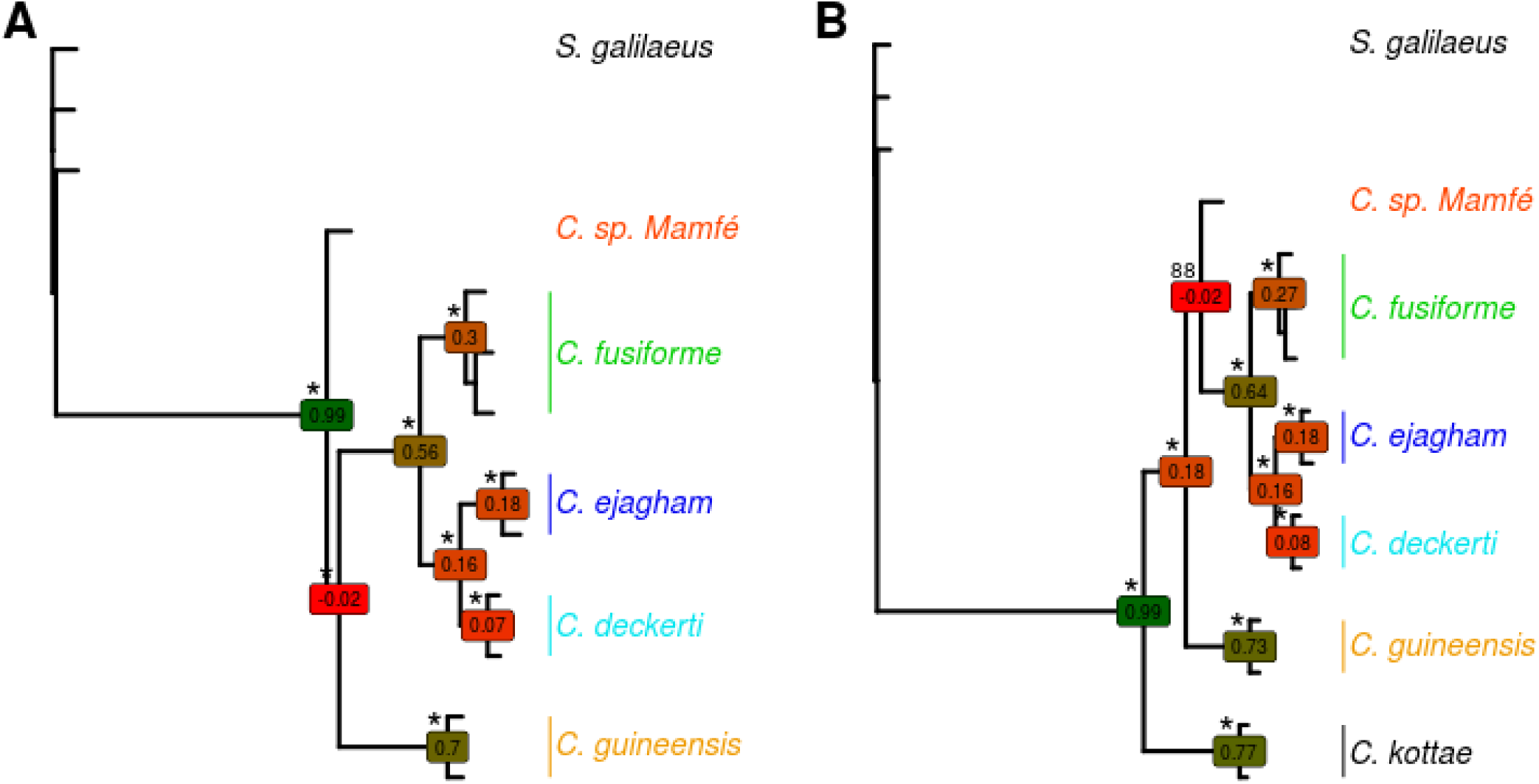
ML trees of concatenated whole-genome sequences with different outgroup configurations. **(A)** Using only *S. galilaeus* as an outgroup; **(B)** Using both *S. galilaeus* and *C. kottae* as outgroups. In both cases, a monophyletic Ejagham *Coptodon* radiation is inferred, as is a sister relationship between *C. deckerti* and *C. ejagham*. However, in (A), *C. guineensis*, and in (B), *C. sp. Mamfé* is inferred to be sister to Ejagham *Coptodon*. Bootstrap support (* = 100% support), and ICA (Internode Confidence All) scores based on ML gene trees for 100kb windows are also shown. In both panels, ICA scores are negative for the node grouping Ejagham Coptodon and the inferred sister species, indicating that the concatenated ML tree does not represent the most common gene tree, which instead has a sister relationship between *C. guineensis* and *C. sp. Mamfé*.

**S2 Fig.**
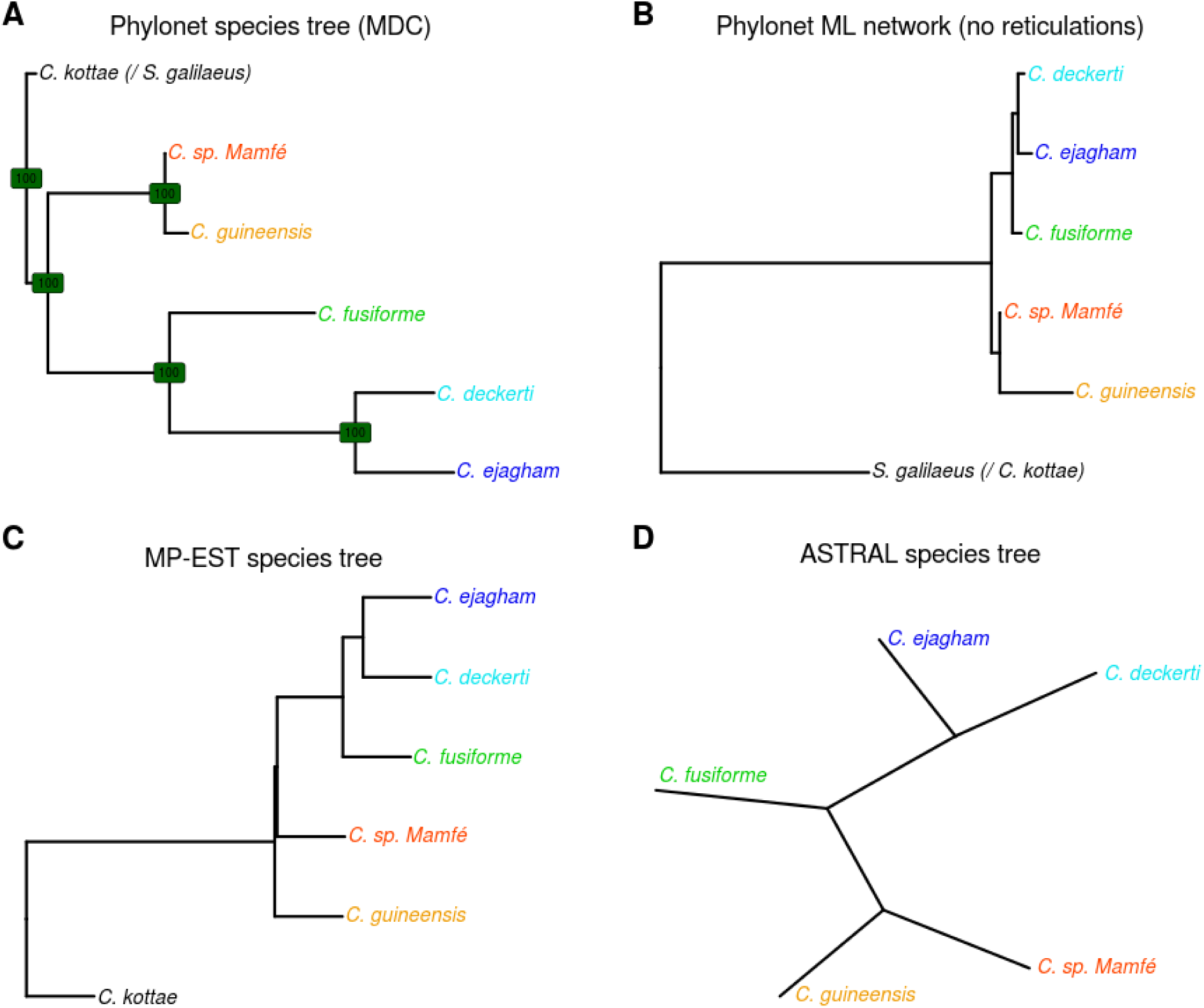
Species trees. All species trees were constructed from gene trees based on 100 kb genomic windows (rooted trees for A-C, and unrooted for D). All species trees have a topology with a monophyletic Lake Ejagham radiation, and a sister relationship between *C. deckerti* and *C. ejagham*. The only difference among the tree topologies is the position of *C. sp. Mamfé*, which is sister to the Ejagham radiation only using the maximum pseudo-likelihood method in MP-EST.

**S3 Fig.**
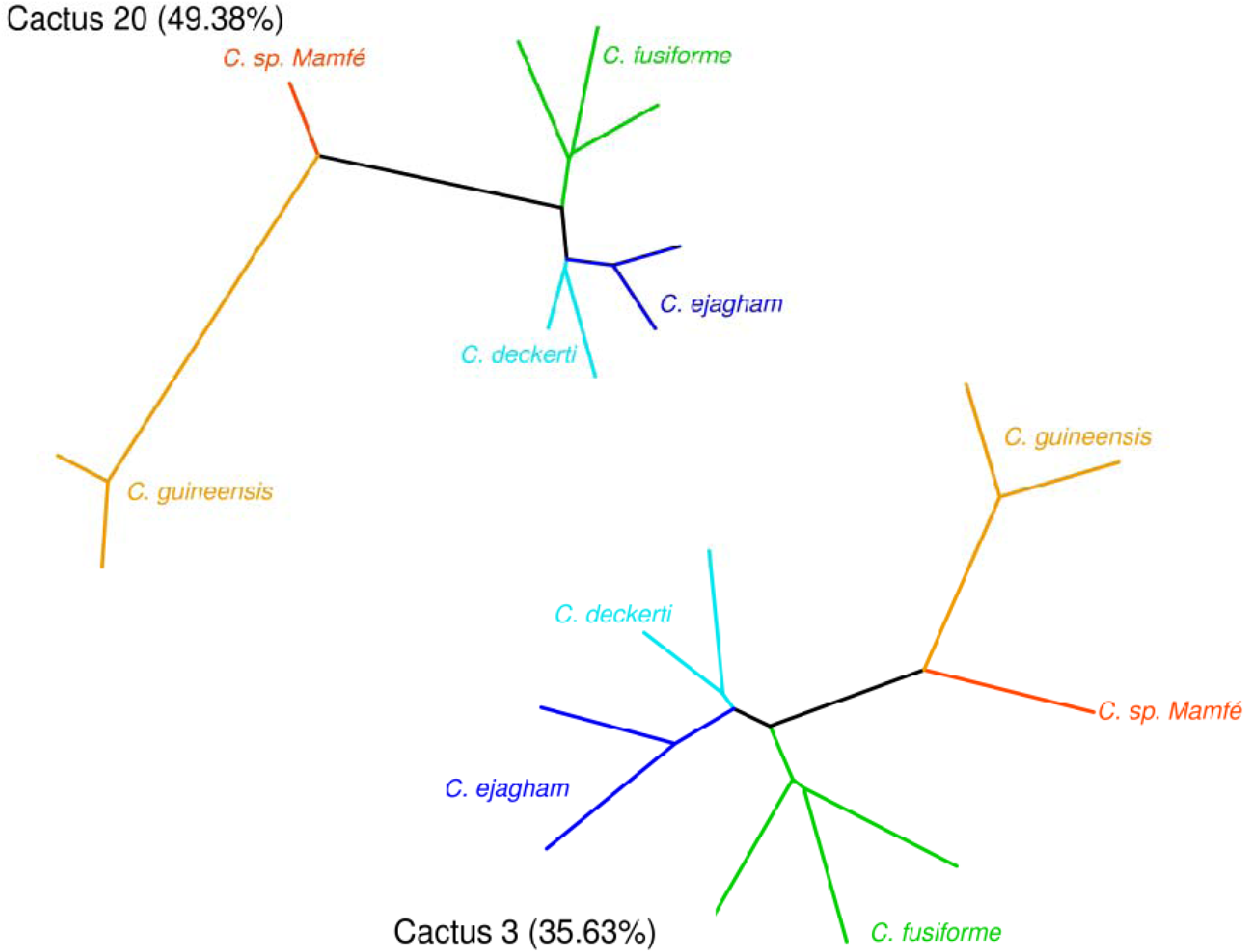
The two most common Saguaro cacti have the same topology with a monophyletic Lake Ejagham radiation.

**S4 Fig.**
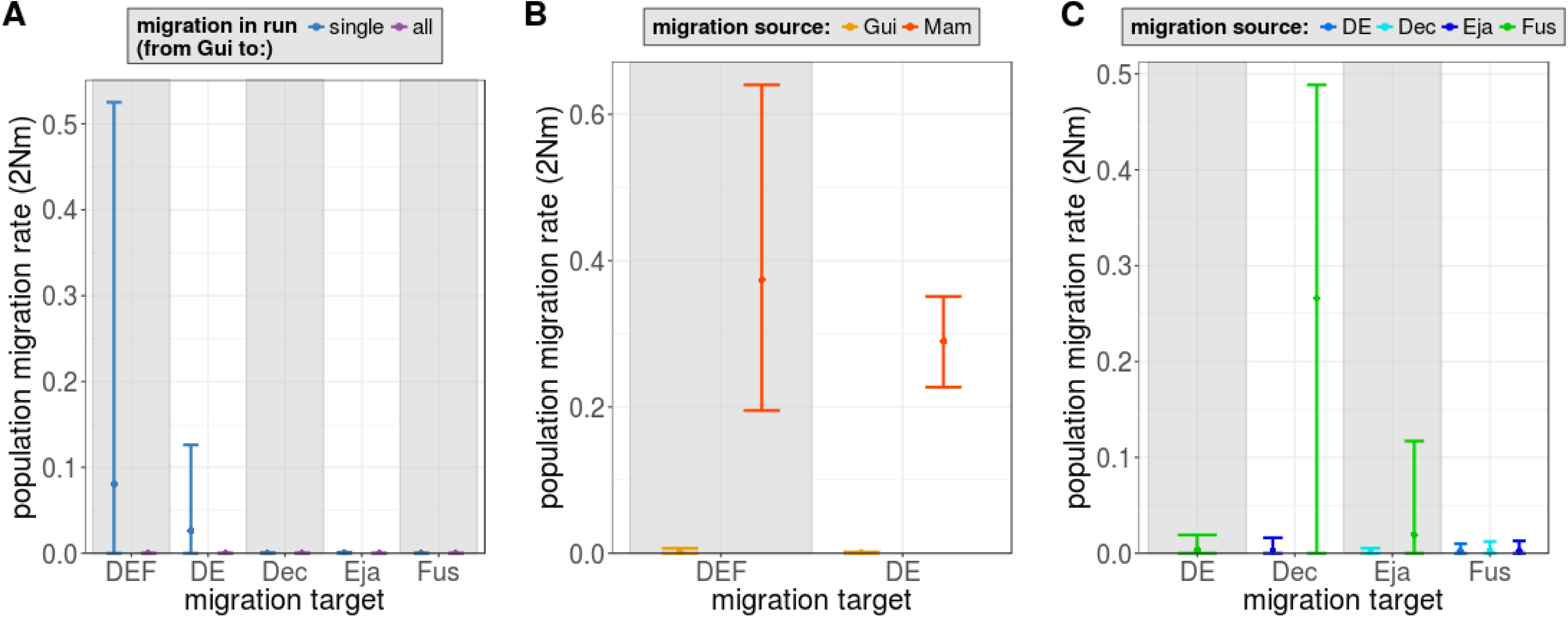
No significant migration from *C. guineensis* or within the Lake Ejagham radiation. Population migration rates estimated by G-PhoCS **(A-B)** from *C. guineensis*, and **(C)** within the Lake Ejagham radiation. In (B), migration bands are estimated simultaneously from *C. sp. Mamfe* and *C. guineensis* to ancestral Lake Ejagham lineages (“DEF”, the ancestor of all thee species, and “DE”, the ancestor of *C. deckerti* and *C. ejagham)*. While in (A), a lot of variance around the migration estimates from *C. guineensis* to DE and DEF are observed, this is no longer the case in (B), when migration from *C. sp. Mamfé* is included. Estimates of migration within the radiation (C-D) are only from runs with single migration bands.

**S5 Fig.**
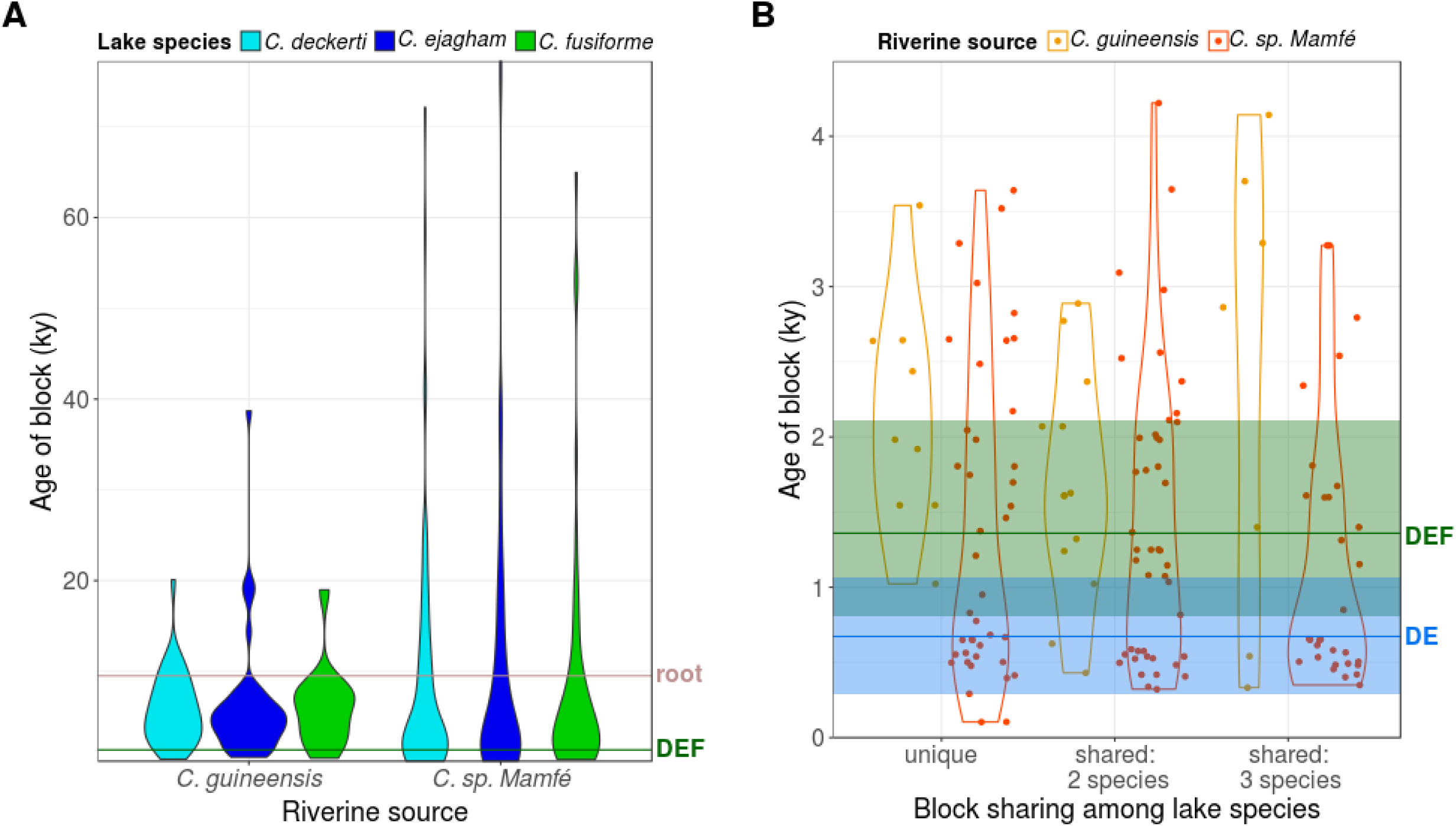
Age distribution of admixture blocks. (**A**) Age distribution of potential admixture blocks prior to filtering by age. The estimated time of divergence between the riverine and lake lineages (line marked with “root”), and the estimated time of the first speciation event within the lake (line marked with “DEF”), as estimated by G-PhoCS, are also shown. Most blocks are estimated to be of more recent origin than the divergence time of the lake lineage, but many others are older and are presumably caused by lineage sorting processes rather than admixture. **(B)** Age distribution of “high-confidence” (age-filtered) admixture blocks by sharing category. Unique blocks do not tend to be younger than shared blocks.

**S6 Fig.**
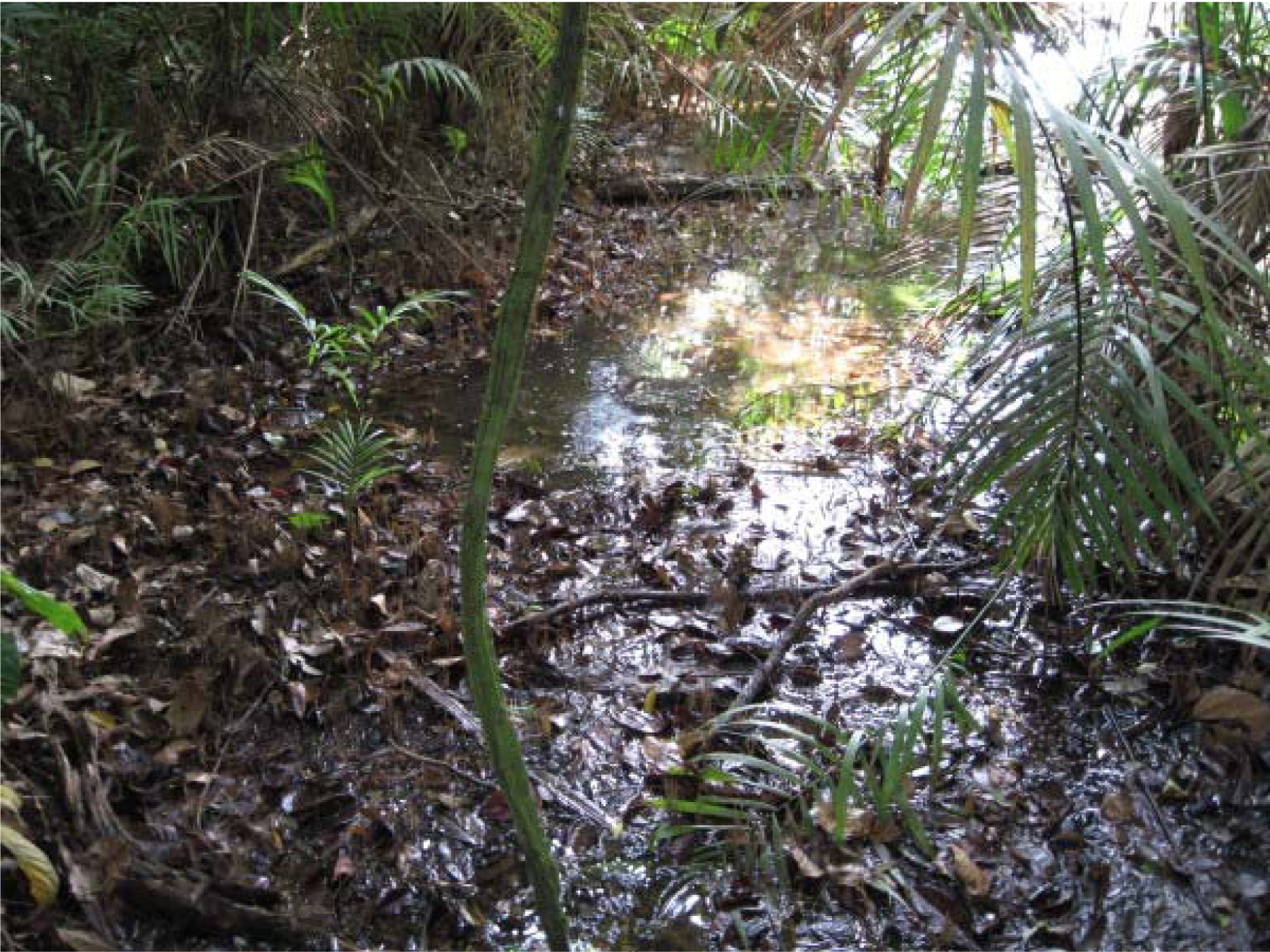
Lake Ejagham’s outlet stream, during the dry season (January 11, 2010).

**S7 Table.** **Monophyly characteristics of each Saguaro cactus.** “Percentage of genome” – The percentage of the genome that is assigned to each cactus. “Radiation monophyletic” – whether (1) or not (0) the Lake Ejagham radiation as a whole forms a monophyletic group to the exclusion of the two riverine species *C. sp. Mamfé* and *C. guineensis*. “Each species monophyletic” – whether (1) or not (0) individuals of each of the three Lake Ejagham are monophyletic.

| cactus<br>nr. | percentage<br>of genome | radiation<br>monophyletic | all species<br>monophyletic |
| --- | --- | --- | --- |
| 0 | 0.56 | 0 | 0 |
| 1 | 2.49 | 1 | 0 |
| 2 | 0.89 | 1 | 1 |
| 3 | **35.63** | **1** | **1** |
| 4 | 0.81 | 0 | 0 |
| 5 | 0.04 | 0 | 0 |
| 6 | 0.89 | 1 | 0 |
| 7 | 1.19 | 0 | 1 |
| 8 | 0.31 | 0 | 0 |
| 9 | 0.29 | 0 | 0 |
| 10 | 0.49 | 0 | 0 |
| 11 | 0.15 | 0 | 0 |
| 12 | 1.12 | 0 | 0 |
| 13 | 0.23 | 0 | 0 |
| 14 | 0.39 | 0 | 0 |
| 15 | 0.25 | 1 | 0 |
| 16 | 0.21 | 0 | 0 |
| 17 | 0.88 | 0 | 0 |
| 18 | 0.10 | 1 | 0 |
| 19 | 0.01 | 0 | 0 |
| 20 | **49.38** | **1** | **1** |
| 21 | 0.06 | 0 | 0 |
| 22 | 0.05 | 1 | 0 |
| 23 | 0.32 | 0 | 1 |
| 24 | 0.01 | 1 | 1 |
| 25 | 0.06 | 1 | 0 |
| 26 | 2.75 | 0 | 0 |
| 27 | 0.15 | 0 | 0 |
| 28 | 0.19 | 1 | 1 |
| 29 | 0.02 | 1 | 0 |
| 30 | 0.06 | 1 | 0 |

**S8 Table.** All species configurations used for calculation of the *f_d_* statistic.

| species A | species B | species C | outgroup |
| --- | --- | --- | --- |
| *C. fusiforme* | *C. deckerti* | *C. sp. Mamfé* | *S. galilaeus* |
| *C. fusiforme* | *C. deckerti* | *C. guineensis* | *S. galilaeus* |
| *C. fusiforme* | *C. ejagham* | *C. sp. Mamfé* | *S. galilaeus* |
| *C. fusiforme* | *C. ejagham* | *C. guineensis* | *S. galilaeus* |
| *C. deckerti* | *C. ejagham* | *C. sp. Mamfé* | *S. galilaeus* |
| *C. deckerti* | *C. ejagham* | *C. guineensis* | *S. galilaeus* |
| *C. deckerti* | *C. fusiforme* | *C. sp. Mamfé* | *S. galilaeus* |
| *C. deckerti* | *C. fusiforme* | *C. guineensis* | *S. galilaeus* |
| *C. ejagham* | *C. fusiforme* | *C. sp. Mamfé* | *S. galilaeus* |
| *C. ejagham* | *C. fusiforme* | *C. guineensis* | *S. galilaeus* |
| *C. ejagham* | *C. deckerti* | *C. sp. Mamfé* | *S. galilaeus* |
| *C. ejagham* | *C. deckerti* | *C. guineensis* | *S. galilaeus* |
| *C. sp. Mamfé* | *C. guineensis* | *C. deckerti* | *S. galilaeus* |
| *C. sp. Mamfé* | *C. guineensis* | *C. ejagham* | *S. galilaeus* |
| *C. sp. Mamfé* | *C. guineensis* | *C. fusiforme* | *S. galilaeus* |
| *C. guineensis* | *C. sp. Mamfe* | *C. deckerti* | *S. galilaeus* |
| *C. guineensis* | *C. sp. Mamfe* | *C. ejagham* | *S. galilaeus* |
| *C. guineensis* | *C. sp. Mamfe* | *C. fusiforme* | *S. galilaeus* |

**S9 Table.**
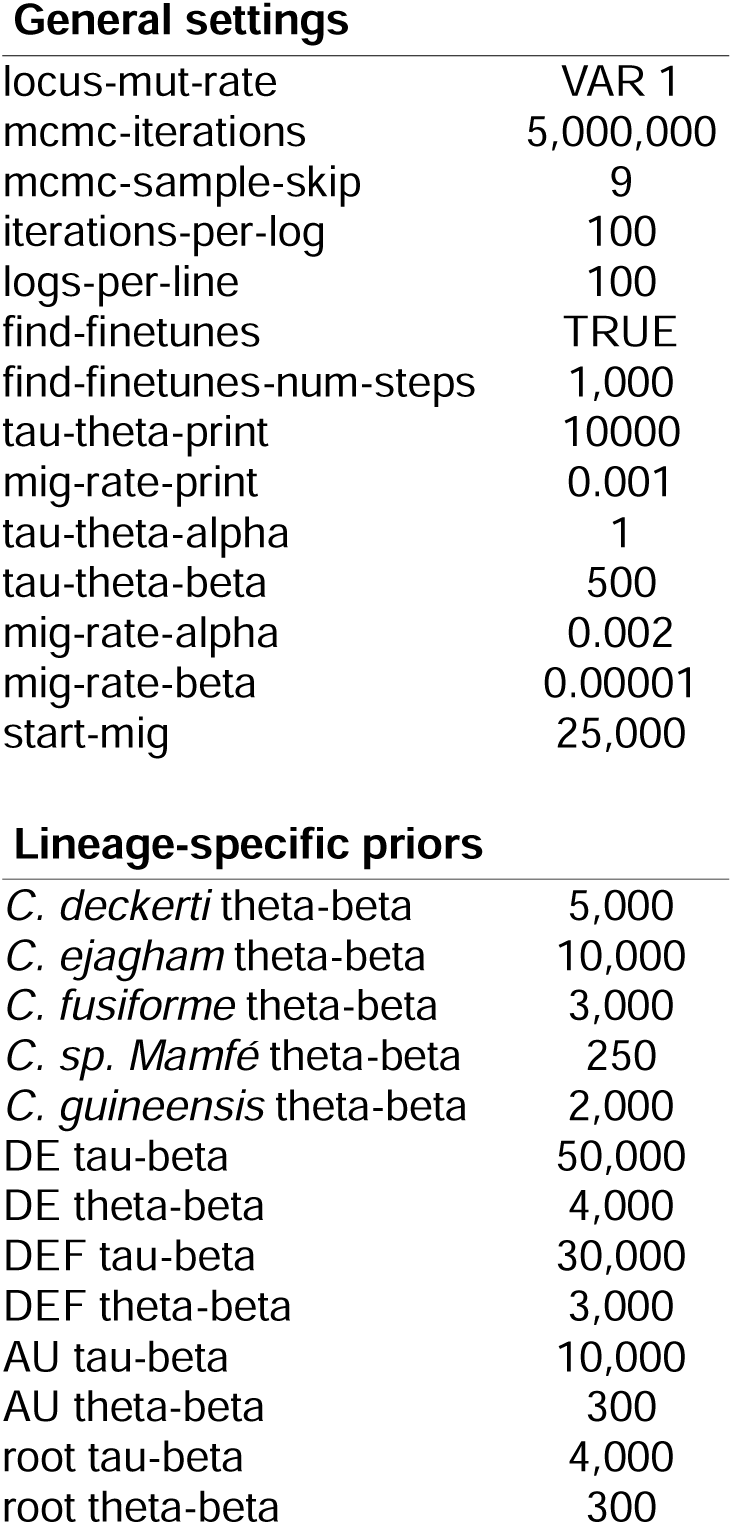
Settings for G-PhoCS.

**General settings**
|  |  |
| --- | --- |
| locus-mut-rate | VAR 1 |
| mcmc-iterations | 5,000,000 |
| mcmc-sample-skip | 9 |
| iterations-per-log | 100 |
| logs-per-line | 100 |
| find-finetunes | TRUE |
| find-finetunes-num-steps | 1,000 |
| tau-theta-print | 10000 |
| mig-rate-print | 0.001 |
| tau-theta-alpha | 1 |
| tau-theta-beta | 500 |
| mig-rate-alpha | 0.002 |
| mig-rate-beta | 0.00001 |
| start-mig | 25,000 |
| **Lineage-specific priors** |  |
| *C. deckerti* theta-beta | 5,000 |
| *C. ejagham* theta-beta | 10,000 |
| *C. fusiforme* theta-beta | 3,000 |
| *C. sp. Mamfé* theta-beta | 250 |
| *C. guineensis* theta-beta | 2,000 |
| DE tau-beta | 50,000 |
| DE theta-beta | 4,000 |
| DEF tau-beta | 30,000 |
| DEF theta-beta | 3,000 |
| AU tau-beta | 10,000 |
| AU theta-beta | 300 |
| root tau-beta | 4,000 |
| root theta-beta | 300 |

**S10 Table.** Olfactory receptor genes found in an admixture block between *C. sp. Mamfé* and *C. deckerti*/*C. Ejagham*.

| Entrez Gene ID | Ensembl Gene ID | Gene description | 1-to-1 orthologues |
| --- | --- | --- | --- |
| 100695228 | ENSONIG00000016134 | olfactory receptor 6N2-like | NA |
| 100694957 | ENSONIG00000016134 | olfactory receptor 2AT4-like | NA |
| 100707735 | ENSONIG00000020687 | olfactory receptor 11A1-like | NA |
| 100692273 | ENSONIG00000020687 | olfactory receptor 6N2-like | NA |
| 100693089 | ENSONIG00000020689 | olfactory receptor-like protein OLF4 | NA |
| 100694162 | ENSONIG00000020690 | olfactory receptor 4C12-like | NA |
| 100691734 | ENSONIG00000020690 | olfactory receptor 13C5-like | NA |
| 100696017 | ENSONIG00000020691 | olfactory receptor 6N2-like | ENSPFOG00000020775 |

